# *SPPiDDRs*: a new gene family in dicot plants involved in DNA-Damage Response

**DOI:** 10.1101/2023.12.20.568739

**Authors:** Valentin Hammoudi, Elisa Goldbecker, Josephine Herbst, Loreen Linnenbrügger, Antje von Schaewen, Reinhard Kunze

## Abstract

Living organisms must maintain the integrity of their genome, and plants are not exempt. In plants, recognition of DNA damage converges at the transcription factor SOG1, a functional homolog of the animal p53 protein. SOG1 directly controls the expression of hundreds of genes and orchestrates a sophisticated network of signaling pathways termed DNA-damage response (DDR). Only recently, several long non-coding RNA (lncRNA) loci were identified to be upregulated by DNA damage, and only a handful have been confirmed to actively contribute to DDR. In this study, we focused on one locus annotated as lncRNA and found that it is strongly and quickly upregulated upon DNA damage and is a direct target of SOG1. Combining *in silico* and experimental analyses, we demonstrate that this locus was wrongly annotated as lncRNA and is in fact a gene coding for a short protein that targets peroxisomes. Consequently, we renamed this locus *<u>S</u>HORT <u>P</u>EROXISOMAL <u>P</u>ROTEIN INDUCED IN <u>D</u>NA-<u>D</u>AMAGE <u>R</u>ESPONSE1* (*SPPiDDR1*). *SPPiDDRs* are well conserved and present in multiple copies across dicot genomes, with Arabidopsis containing two additional copies, *SPPiDDR2* and *SPPiDDR3*. The *AtSPPiDDR* paralogs differ on the transcriptional level, *SPPiDDR3* being the least active. *SPPiDDR1* and *SPPiDDR2* are both also induced by salt, a stress treatment known to indirectly induce DNA damage via oxidative stress. We show that these two genes act redundantly and inhibit plant growth in response to salt stress.

## INTRODUCTION

DNA integrity can be altered by a plethora of factors, such as ionizing radiation (X-rays and γ-irradiation [γ-IR]), UV-light, metals (*e.g.* aluminum), or chemicals (*e.g.* zeocin, bleomycin, zebularine)^1^. In addition, Reactive Oxygen Species (ROS) may also affect DNA integrity, meaning that metabolism or conditions that cause oxidative bursts, like salt stress, can also indirectly damage DNA^2^. These genomic alterations can manifest in different ways: base mismatches, pyrimidine dimers, intra- or inter-strand crosslinks, single-strand breaks (SSB), or double-strand breaks (DSB). When such DNA lesions occur, they must be repaired quickly and efficiently before the next cell division starts, in order to prevent mutations from being carried over into the progeny.

Once DNA damage is detected in eukaryotic cells, signaling cascades are initiated by two serine/threonine kinases: ATAXIA-TELANGIECTASIA MUTATED (ATM) and ATAXIA TELANGIECTASIA-MUTATED AND RAD3-RELATED (ATR)^3–5^. In mammals, ATM and ATR indirectly activate p53, also known as the “guardian of the genome”, which is the transcription factor (TF) that mounts and orchestrates the DNA-damage response (DDR)^6,7^ Strikingly, plants lack p53, but ATM directly phosphorylates and activates the plant-specific TF SUPPRESSOR OF GAMMA IRRADIATION1 (SOG1), which is the plant master regulator of DDR. It is assumed that ATR operates similarly, even though SOG1 phosphorylation by ATR has been shown only *in vitro*^8–10^. SOG1 belongs to the NAC [NO APICAL MERISTEM (**N**AM), ARABIDOPSIS TRANSCRIPTION ACTIVATION FACTOR (**A**TAF), CUPLSHAPED COTYLEDON (**C**UC)] TF family, known to be involved in stem cell maintenance, senescence, and (a)biotic stress response(s)^9,11–14^. Despite the absence of sequence homology, SOG1 is the functional homolog of p53, since both TFs act on functionally equivalent target genes in plants and animals^15–17^. Upon activation, both SOG1 and p53 trigger a massive transcriptional reprogramming that results in cell-cycle arrest, DNA repair, and in some cases also in programmed cell death or endoreduplication.

While up to now more than 3000 target genes of p53 were reported^18,19^, efforts to identify direct SOG1 target genes on a large scale started only recently by comparing transcriptomes of wild-type plants and *sog1* mutants, thus confirming *bona fide* direct targets of SOG1 by Chromatin-Immunoprecipitation sequencing (ChIP-seq)^15,17,20,21^. More than 300 direct SOG1 target genes containing the palindromic CTT(N)_7_AAG consensus SOG1-binding motif have been discovered with this approach since then^15^. Besides cell cycle (e.g. *CYCB1* and *SMR7*), DNA repair (e.g. *BRCA1*), and programmed cell death (e.g. *PLA2A*), SOG1 direct target genes were found to be involved in processes like transcriptional and posttranscriptional regulation, oxidative stress and defense responses, uncovering new facets of the DDR in plants^9,15,17,20^.

In mammals, p53 also directly controls the expression of long non-coding RNA (lncRNA)^22–24^. The term lncRNA refers to transcripts that present two key features: length > 200 bases and low likelihood of being translated, the latter being suggested by either the absence of an Open Reading Frame (ORF), poor protein-coding potential, no ribosomal binding detected via ribosome profiling, and/or absence of matching peptides via proteomics, despite transcriptional activity^25^. While the involvement of lncRNAs in DDR and their transcriptional control by p53 is well established in mammals^22,26,27^, the role of lncRNAs in plant DDR has emerged only recently. We previously identified 56 lncRNAs whose expression is induced in *Arabidopsis thaliana* seedlings three hours after an irradiation with 80 Gray X-rays^28^. The recorded expression change of 93% of these lncRNA was ATM-dependent, implying a potential role in DDR. More recently, we focused on one of these lncRNAs, *LINDA*, and found that it is involved in the accurate execution of the DDR in Arabidopsis roots^29^. Not only is *LINDA* needed for the regulation of its flanking gene, but also for the fine-tuning of the DDR after the occurrence of DNA double-strand breaks. Another study confirmed the involvement of *LINDA* in DDR and studied three additional lncRNAs in Arabidopsis^30^. One of these studied lncRNAs, annotated on Araport11 as a novel transcribed region (*AT4G09215*), was strongly upregulated by UV-C irradiation. It also corresponds to the lncRNA *RLFS_026432*, which was one of the most strongly induced lncRNAs upon X-ray exposure in our previous study (more than 225 fold-change), of note partially in an ATM-dependent manner (see Fig. 3A of Wang et al., 2016)^28^.

In this study, we investigated the *RLFS_026432* locus further in the context of DDR. We found that *RLFS_026432* was mis-annotated as lncRNA, and is in fact a gene encoding a short uncharacterized protein. We revealed that genes with high sequence identity are present in multiple copies across dicot genomes, Arabidopsis possessing two additional ones. *RLFS_026432* is strongly induced by diverse kinds of DNA stressors and is a direct target of SOG1, supporting the hypothesis that this locus plays a pivotal role in DDR. We found that *RLFS_026432* and one of its paralogs contribute redundantly in the response to salt stress, a condition that indirectly triggers DNA damage via reactive oxygen species (ROS). Finally, we showed that one of the paralogs is also active in normal growth conditions, suggesting that the functions of this newly discovered plant gene family extend beyond merely stress responses.

## RESULTS

### RLSF_026432 is induced in response to DNA damage

*RLFS_026432* was found to be one of the most highly induced lncRNAs upon DNA damage caused by X-rays or UV-C^28,30^. To determine the optimal conditions for its induction by UV-C, different doses were tested and expression of *RLFS_026432* was measured after 3 hours, the same time frame as in Wang et al. (2016)^28^. A UV-C dose of 3 kJ/m² led to the strongest induction of *RLFS_026432* (**Fig. 1A**). Already 30 minutes after UV-C exposure, *RLFS_026432* induction was detectable, increased after 1 hour (142 fold-change) and peaked at 3 hours (205 fold-change) before progressively decreasing and persisting up to 72 hours (37 fold-change) (**Fig. 1A**). Consequently, we selected 3 hours after a 3 kJ/m² dose of UV-C as the optimal sampling time point to investigate the transcriptional activation of this locus.

**Figure 1:**
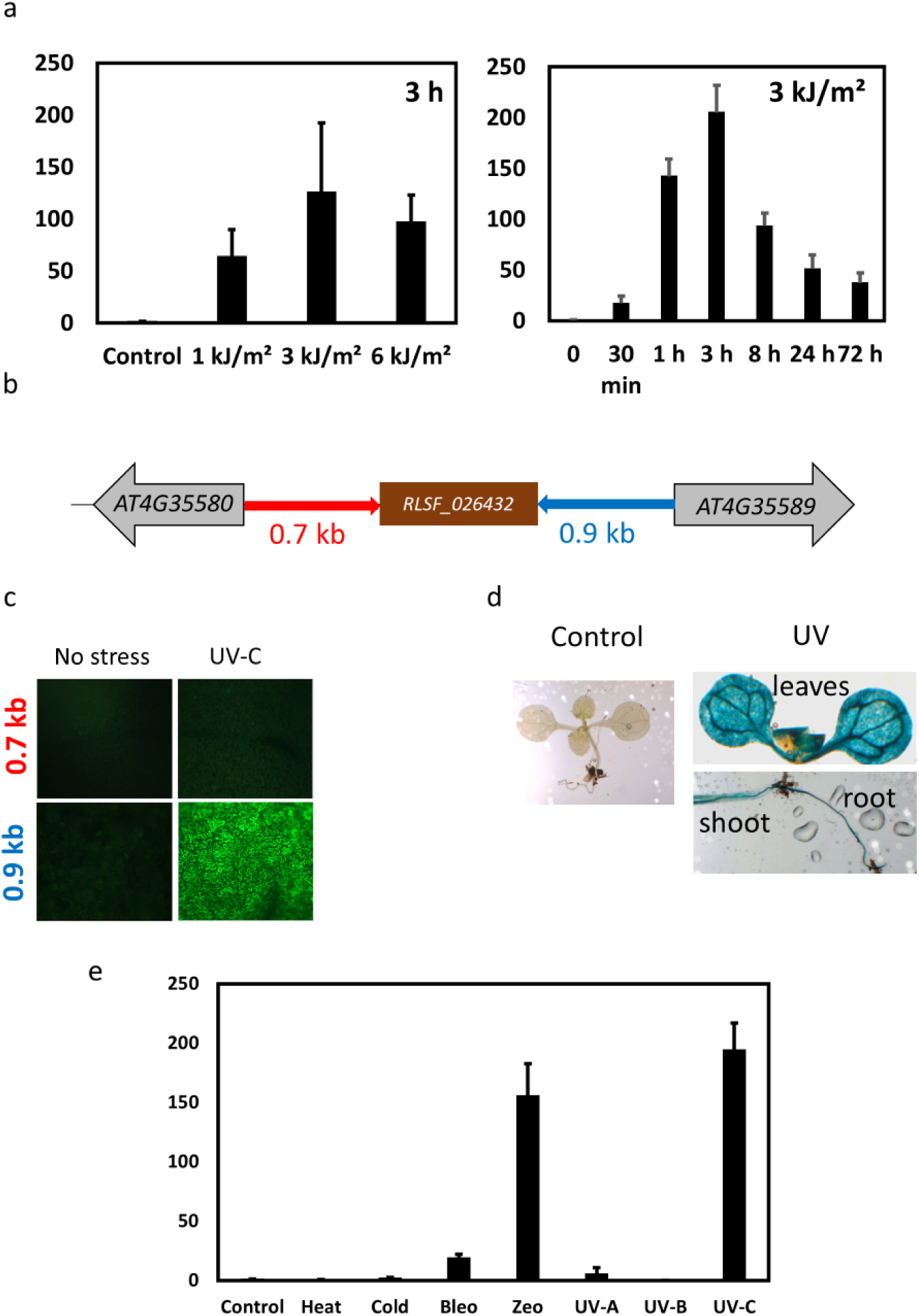
*RLSF_026432* is induced in response to DNA damage. a- Transcription analysis by RT-qPCR of *RLFS_026432* in response to UV-C exposure, either 3 hours after different doses or at different time points with a 3 kJ/m² dose. Data were normalized to the values in non-treated wild-type plants (Control = 1). b- Schematic representation (not to scale) of the genetic environment of *RLSF_026432.* The 728 bases (0.7 kb) separating *RLSF_026432* from *AT4G35580*, and the 975 bases (0.9 kb) separating *RLSF_026432* from *AT4G35589* (in red and blue, respectively, with the arrow indicating the 5’-3’ orientation) were both cloned to get two *promoter::GFP-GUS* cassettes reporting on their respective promoter activity. c- 3 to 4-week old seedlings were exposed to UV-C and leaf discs were analyzed by epifluorescence microscopy. d- 3-week old seedlings carrying the *proRLSF_026432::GFP-GUS* cassette were exposed to UV-C and after 3 hours GUS-stained. e- Transcription analysis by RT-qPCR of *RLFS_026432* in response to several stresses: heat (35°C for 3 hour), cold (4°C for 3 hours), bleo (3 hours in bleomycin 1 μg/ml), zeo (6 hours in zeocin 40 µg/ml), UV-C (3 kJ/m²), UV-B (2 kJ/m²) and UV-A (2 kJ/m²). Data were normalized to the values in non-treated wild-type plants.

To determine the gene orientation of *RLFS_026432*, we cloned the intergenic sequences separating the locus from its neighboring genes (*AT4G35580* and *AT4G35589,* respectively), upstream of a *GFP-GUS* fusion coding sequence and introduced them into the Arabidopsis genome via stable transformation (**Fig. 1B**). T2 homozygous plants were then irradiated with a 3 kJ/m² UV-C dose. While the 0.7 kb sequence between *AT4G35580* and *RLFS_026432* had no "forward" promoter activity, the 0.9 kb sequence between *AT4G35589* and *RLFS_026432* exhibited strong "reverse" promoter activity, demonstrating that the *AT4G09215* locus is transcribed from the reverse strand of the Arabidopsis genome, consistent with the Araport11 annotation (**Fig. 1C**). This 0.9 kb intergenic region therefore contains the UV-C inducible *RLFS_026432* promoter. We then determined the tissues in which *RLFS_026432* is expressed using GUS staining of seedlings (**Fig. 1D**). No expression of *RLFS_026432* was observed in basal conditions, while a strong induction was detected in leaves, shoots, and roots after UV-C irradiation. Overall, our data indicate that the expression of *RLFS_026432* induced by UV-C is quick, strong, and systemic.

To determine whether *RLFS_026432* induction is also triggered by other DNA-damaging conditions, we treated seedlings with bleomycin or zeocin, two chemicals inducing DNA double-strand breaks. Both chemicals induced *RLFS_026432* expression. With zeocin, the induction was comparable to the one observed with UV-C and much stronger than with bleomycin (>150 fold-change for zeocin, >20 fold-change for bleomycin) (**Fig. 1E**). In addition, we examined *RLFS_026432* with RNA-seq data from Bourbousse et al. (2018). While we found no read matching this locus in Arabidopsis seedlings under normal conditions, γ-IR led to an induction comparable to DDR genes such as *SMR7* or *BRCA1*^17^. Furthermore, we tested UV-A and UV-B irradiation at a dose comparable to UV-C, but none led to an increase of lncRNA expression (**Fig. 1E**). UV-A and UV-B are known to also create DNA lesions, so these results indicate that not all DNA-damage stressors induce *RLFS_026432*. Finally, we tested two conditions that do not affect DNA integrity, heat and cold, but none led to a transcriptional activation of *RLFS_026432*. In summary, *RLFS_026432* was found to be induced by X-rays^28^, UV-C^30^ (and this study), zeocin^30^ (and this study), bleomycin, and γ-IR^17^, all being strong DNA-damage stressors.

### SOG1 directly controls RLSF_026432 expression

SOG1, the master regulator of DDR in plants, was recently shown to control the expression of the lncRNA *LINDA*, which is involved in the DDR in Arabidopsis^29^. Thus, we investigated whether *RLFS_026432* may be a direct target of SOG1 too. While RNA-seq data from Bourbousse et al. (2018) showed an induction of the *RLFS_026432* locus in wild-type plants treated with γ-IR, transcriptional induction was fully abolished in the *sog1* mutant (**Fig. 2A)**. This indicates that *RLFS_026432* belongs to the SOG1-pathway. In line with this, we identified a SOG1-binding motif (CTT(N)_7_AAG) in the promoter of the lncRNA. Examining ChIP-seq data with flagged SOG1 from Bourbousse et al. (2018), we found that SOG1 indeed binds the promoter of *RLFS_026432*, exactly at the location of the palindromic CTT(N)_7_AAG motif. Taken together, these data show that *RLSF_026432* is a direct target of SOG1.

**Figure 2:**
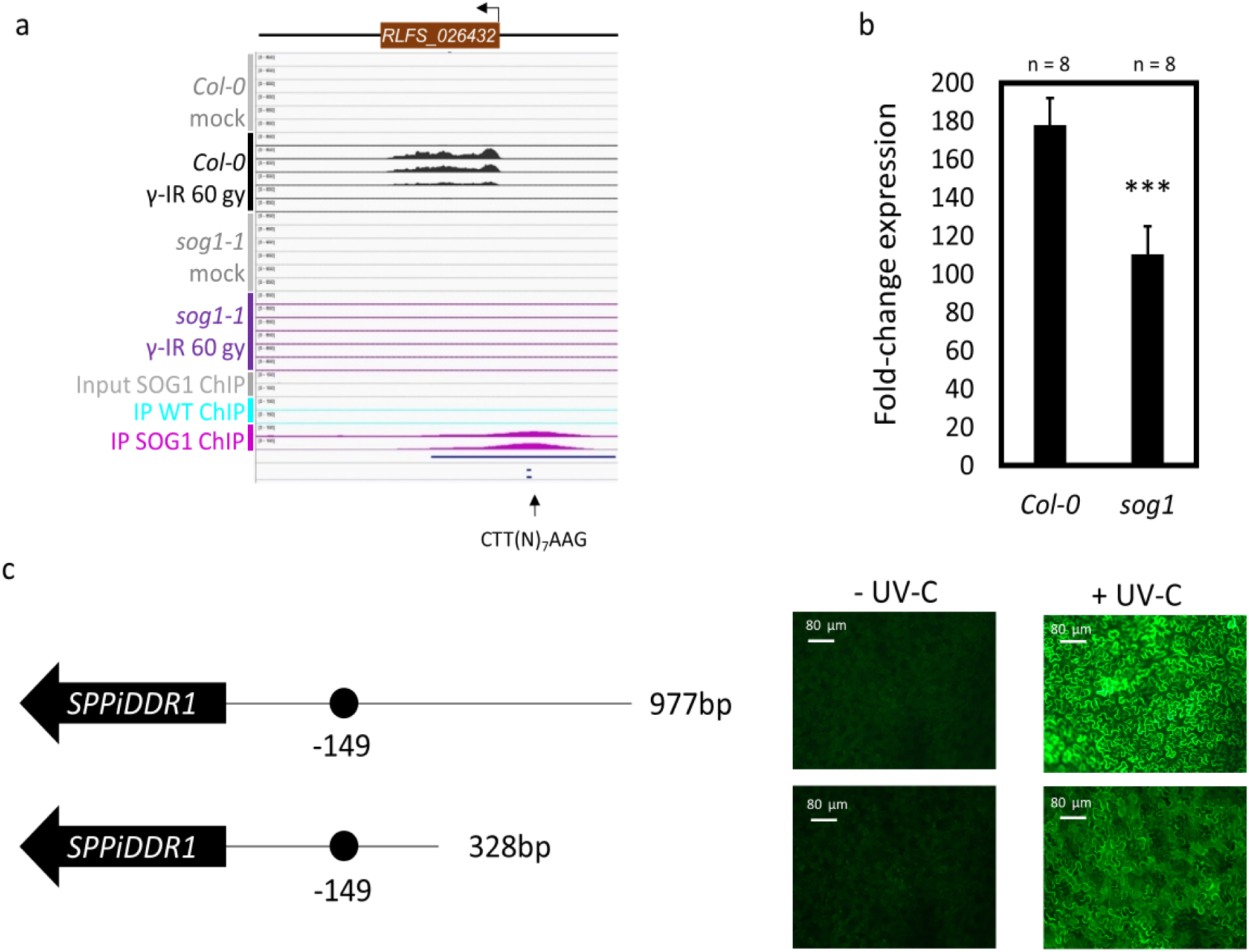
*RLSF_026432* is a direct target of SOG1. a- Depiction of reads from RNA-seq and SOG1 ChIP-seq datasets from Bourbousse et al., 2018 on the *RLFS_026432* locus. Each RNA-seq block (Col-0 or *sog1*, after exposure or not (mock) to γ-IR (60 grays)) contains 6 tracks that correspond to 6 different time points: 0, 1.5 hour, 3 hours, 6 hours, 12 hours and 24 hours after the exposure to γ-IR. ChIP-seq blocks correspond to reads obtained from plants carrying a *pSOG1:SOG1-3xFLAG* construct as bait, compared to their respective input and wild-type controls. Each block is composed of 2 tracks that correspond to 2 replicates of samples picked 1 hour after exposure to γ-IR (60 grays). Scales are identical in between RNA-seq reads, and in between ChIP-seq reads. The CTT(N)_7_AAG motif was found manually. b- Expression analysis with RT-qPCR of *RLSF_026432* in wild-types and in *sog1* mutants treated with UV-C, normalized to expression level in non-treated plants. Three stars indicate a statistical difference with P-value < 0.001. c- Representation of *RLSF_026432* and its promoter. Full-length promoter is composed of the 975 bases upstream the transcript, truncated promoter is composed of only the 326 most proximal base of the full-length promoter. Black dots are SOG1 binding sites. Arabidopsis carrying each of these constructs were analyzed by epifluorescence microscopy before and after UV-C exposure.

To verify that *RLSF_026432* regulation by SOG1 is specific to γ-IR, we analyzed the expression of the locus in *sog1* mutant plants exposed to UV-C. The induction was impaired, indicating that SOG1 is involved in the transcriptional activation of *RLFS_026432* also upon UV-C irradiation (**Fig. 2B**). However, transcription was not fully abolished, like when using γ-IR, suggesting that (an)other TF(s) probably control(s) *RLFS_026432* expression. We cloned a truncated version of the *RLFS_026432* promoter into the *GFP-GUS* expression reporter cassette, consisting of the 328 most proximal bases that still contain the binding site of SOG1, and then generated transgenic Arabidopsis lines (**Fig. 2C**). Following UV-C irradiation, a GFP-fluorescence signal was still observed with the truncated promoter, albeit weaker than with the full-length promoter (**Fig. 2C**). Taken together, these data indicate that *RLFS_026432* is directly controlled by SOG1, even though other TFs may also be involved in its transcriptional regulation, at least in response to UV-C. Since most direct SOG1 targets are involved in the DDR, it is very likely that *RLFS_026432* contributes to this pathway.

### RLFS_026432 encodes a Short Peroxisomal Protein induced in DDR

Although *RLFS_026432* was annotated as “lncRNA” in the Plant Long noncoding RNA Database^31,32^ and as “novel transcribed region” in TAIR10^33^ (*AT4G09215*), an ORF search revealed the presence of four short ORF, from here on referred to as ORF1-4 (**Fig. 3A**). ORF3 is the longest with 59 amino acids, in the same orientation as *RLFS_026432*, and starts at the 5’ end of the locus. Interestingly, a nucleotide BLAST search with ORF3 yielded homologous sequences from *Arabidopsis thaliana* but also from *Arabidopsis lyrata* and *Arabidopsis arenosa* (**Suppl. Table S1**). Furthermore, a protein BLAST search with ORF3 revealed homologous sequences from various *Brassicaceae* species, indicating a more stringent conservation at the protein level than at the DNA level (**Suppl. Table S1**). All hits were approximately 60 amino-acid long and annotated as “hypothetical” and/or “uncharacterized” proteins. An alignment of seven of the putative *Brassicaceae* ORF3 orthologs confirmed the strong conservation, suggesting that they are related (**Fig. 3B**). An alignment of the genomic DNA of these loci showed that conservation is confined to nucleotides between the potential start and stop codon of ORF3 (**Fig. 3C**). In addition, the SOG1-binding site also showed strong conservation. Obviously, a selection pressure applies to these potential coding sequences as well as the promoter elements that control transcriptional activation upon DNA-damage stress.

**Figure 3:**
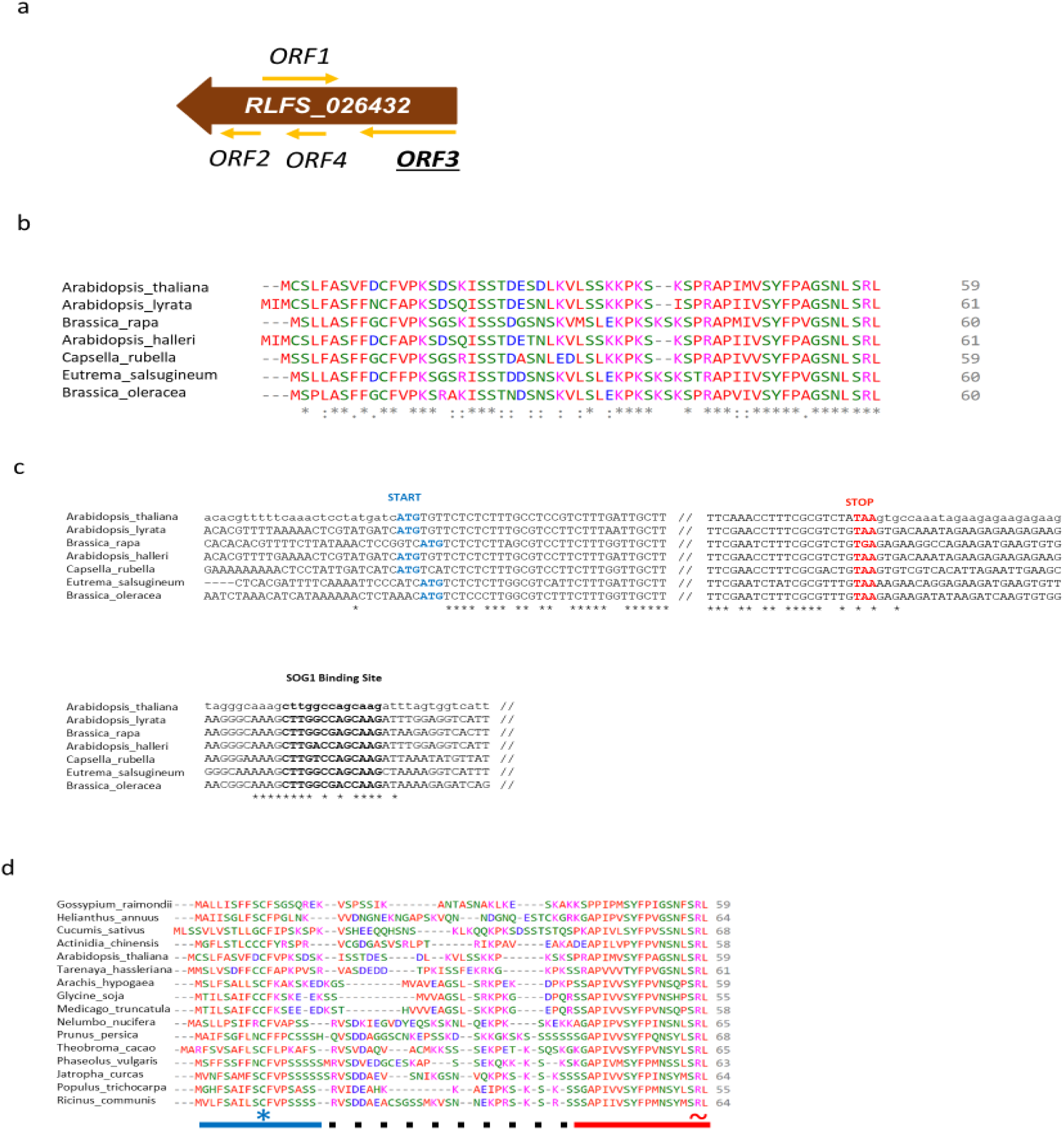
A strong selection pressure applied to the conservation of an ORF in *RLSF_026432* across eudicots. a- Schematic representation of the 4 ORFs found in *RLSF_026432* using “ORF finder” on NCBI. b- Protein alignment of 7 Brassicaceae hits obtained with a protein BLAST with ORF3. The protein BLAST search with ORF3 yielded several hits for each species. For clarity, a single hit per species was used for the protein alignment. The red line indicates the most conserved region. Stars indicate strict conservation of the amino acid in all the sequences shown, colons indicate conservation of amino acids with strongly similar properties, periods indicate conservation of amino acids with weakly similar properties. c- gDNA alignment of the 7 Brassicaceae *OFR3* sequences from (b) at the beginning and. the end of the potential coding sequence, as well as at the SOG1 binding site identified in *RLSF_026432* promoter. Stars indicate strict conservation of the base in all the sequences shown. d- Protein alignment of hits obtained from a protein BLAST search with the last 15 residues of Arabidopsis RLSF_026432 ORF3. For each species, the BLAST yielded several sequences, only the hit with the best score was selected for the protein alignment for clarity. The blue line indicates the conserved N-terminal end with namely a strictly conserved Cystein (market with *), the red line indicates the conserved C-terminal end with a strictly conserved SRL motif (marked with ∼) and the dotted line indicates the variable central area.

Of note, the last fifteen C-terminal amino acids appeared to be particularly conserved (**Fig. 3B**). This short sequence was used for another protein BLAST, yielding many new homologous sequences. The full-length sequences of the top 100 hits were all annotated as “uncharacterized potential peptides”, composed of around 60 amino acids that belong to the *Brassicaceae* but also other clades of dicots (**Suppl. Table S1**). Reiterating the same protein BLAST, excluding the *Brassicaceae*, yielded more sequences from additional dicots, with all sequences displaying the same characteristics (**Suppl. Table S1**). By reiterating the BLAST search, restricting it to individual plant species, we identified related sequences from various dicots, including many crops like *Glycine soja, Helianthus annuus, Theobroma cacao*, and *Prunus persica* (**Fig. 3D**). All sequences displayed three distinct features: (i) a particularly high serine content, reaching 24% in *Arabidopsis thaliana* and 29% in *Prunus persica*, presenting sometimes in the form of contiguous repeats (*e.g.* six serines in *Jatropha curcas* and seven in *Prunus persica*); (ii) high similarity of the N-terminal end, with a strictly conserved cysteine (position 11 in *Arabidopsis thaliana*), and (iii) high conservation of the C-terminal end, especially the last three residues (SRL). Taken together, these data suggest that *RLFS_026432* is a gene conserved in dicots that encodes a short protein rather than a lncRNA.

To obtain experimental evidence that *RLFS_026432* truly represents a coding sequence, we searched for peptides matching its potential amino acid sequence in proteomic datasets, but in vain. Yet, ribosome-profiling datasets revealed that ribosomes are binding to this locus, which is another hint that *RLFS_026432* is most likely translated (GWIPS-viz online server; **Suppl. Fig. S1**). To experimentally determine whether *ORF3* codes for a short protein, an expression vector consisting of the native *RLFS_026432* promoter and *ORF3*, lacking its stop codon, was fused to *GFP* without start codon (*proRLFS_026432::ORF3-GFP)* and Agro-infiltrated into *Nicotiana benthamiana* leaves (**Fig. 4A**). The *proRLFS_026432::ORF3* sequence of this expression cassette corresponds to the genomic DNA block of the locus, driven by its native promoter. If *ORF3* is translated, the translation will initiate at the start codon of *ORF3* and yield ORF3-GFP fusion proteins. If *ORF3* is not translated, translation of this *ORF* can start only from the next start codon, which is in the center of *GFP*, yielding only a truncated protein that does not fluoresce. After the construct was transiently introduced into *N. benthamiana* leaves, no GFP signal was observed under control conditions. In contrast, a strong fluorescence signal was observed in leaves irradiated with UV-C (**Fig. 4B**). This unambiguously demonstrates that *RLFS_026432* codes for a protein and that this locus was wrongly annotated as a lncRNA/novel transcribed region.

**Figure 4:**
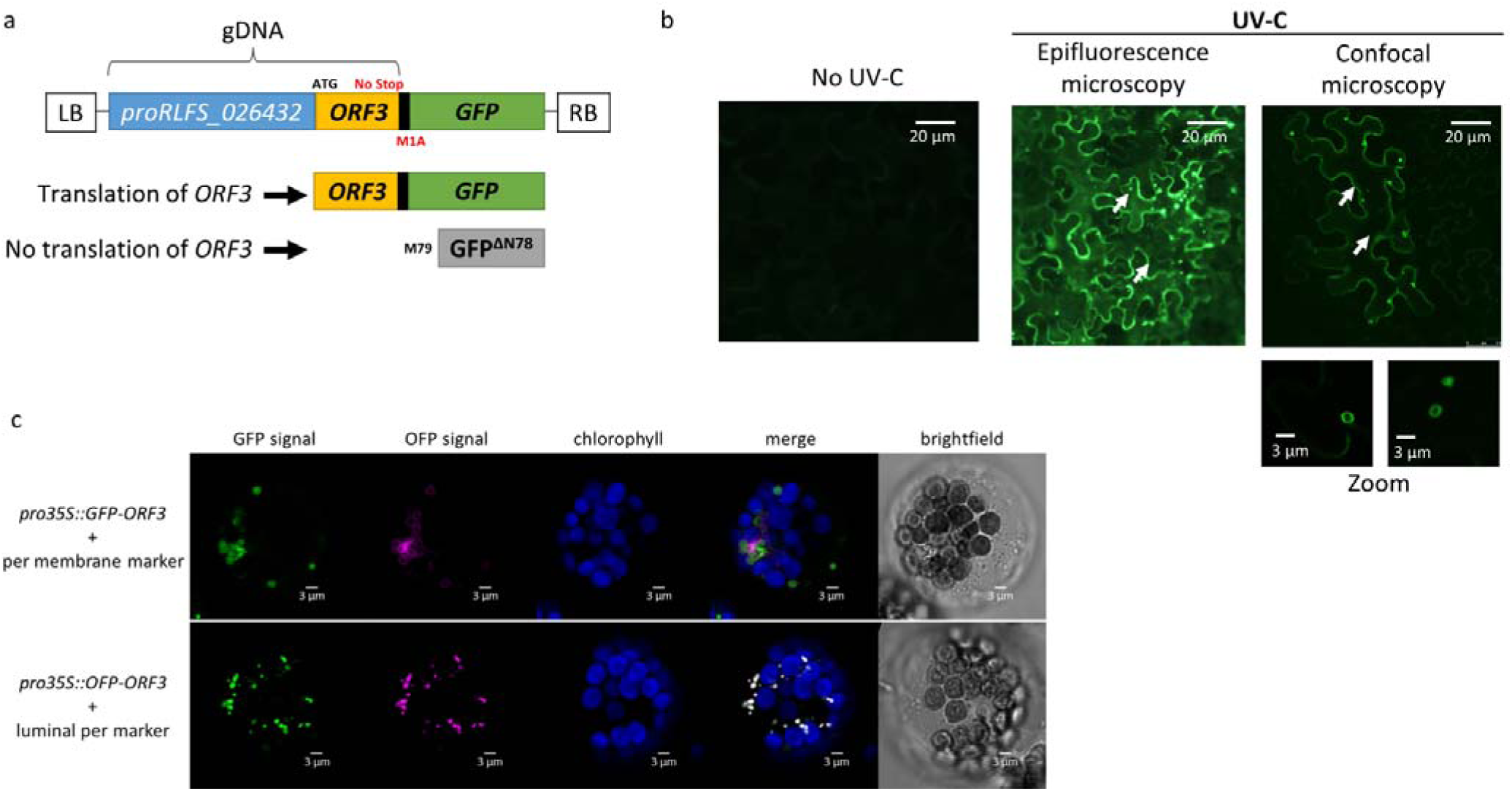
*RLFS_026432* OFR3 encodes a peroxisomal protein. a- Representation of the expression cassette *proRLFS_026432::ORF3-GFP* for transient expression in tobacco leaves. The *proRLFS_026432::ORF3* fragment corresponds to 1158 bases from Arabidopsis gDNA. The removal of the stop codon from ORF3 and of the start codon from the GPF makes it a translation reporter system. After transcriptional activation, if ORF3 undergoes translation, an ORF3-GFP fusion protein will be produced. If ORF3 does not undergo translation, the translation will not start at the start codon of ORF3. As the start codon of GFP has been deleted, the next potential start codon in frame is in position 79 of GFP, which yields a truncated non-fluorescent GFP. b- *proRLFS_026432::ORF3-GFP* were transiently expressed in tobacco leaves. 2 days after infiltration, leaves were exposed to UV-C, and 20 to 24 hours later, the presence of GFP fluorescence was assessed, with epifluorescence microscopy first, followed by confocal microscopy. White arrows indicate subcellular compartments revealed by the GFP signal. c- Arabidopsis protoplasts were cotransformed with a vector carrying a *35S::GFP-ORF3* cassette and a *35S::OFP-PEX16* (top) or a *35S::OFP-PGL3_C-short-SRL* (bottom) cassette, PEX16 and *PGL3_C-short-SRL* being a marker of peroxisomal membrane. For (b) and (c), white bars indicate the scales.

Not only did the detectable fluorescence overlapped with the typical shape of epidermal cells, which suggests that the GFP signal originates from the cytoplasm, the plasma membrane, the cell wall or the apoplast, but also from the periphery of 2 - 3 µm wide subcellular compartments (**Fig. 4B**). In fact, the conserved C-terminal SRL motif, found in all ORF3 sequences, corresponds to a Peroxisomal Targeting Signal of type 1 (PTS1) (**Fig. 3B** and **3D**). Such a motif is recognized by PEROXIN5 (PEX5), which directs proteins containing the PTS1 to peroxisomes^34^. Co-transformation of Arabidopsis protoplasts with *pro35S::GFP-ORF3* and *pro35S::OFP-PEX16* or *pro35S::OFP-PGL3_C-short-SRL* (two markers for the peroxisome membrane or lumen, respectively^35^) showed that ORF3 proteins localize within peroxisomes (**Fig. 4C**). Consequently, *RLFS_026432 ORF3* was renamed *<u>S</u>HORT <u>P</u>EROXISOMAL <u>P</u>ROTEIN INDUCED IN <u>D</u>DR<u>1</u>* (*SPPiDDR1*). Remarkably, the two experiments did not result in the same localization: while *proRLFS_026432::ORF3-GFP* localized at the periphery of peroxisomes (**Fig. 4B**), *pro35S::GFP-ORF3* was detected in the peroxisome matrix (**Fig. 4C**). To test if GFP fused to the C-terminal end of SPPiDDR1 impairs proper localization (disturbing functionality of the PTS1), we repeated transient expression in *N. benthamiana* leaves with a *proRLFS_026432::mCherry-SPPiDDR1* construct, but still obtained a pattern corresponding to localization at the peroxisome periphery upon UV-C exposure (**Suppl. Fig. S2**). Considering that (i) the *pro35S::GFP-ORF3* (newly *pro35S::GFP-SPPiDDR1)* construct without UV-C irradiation resulted in fluorescence of the peroxisome lumen, (ii) an impairment of GFP-SPPiDDR1 localization caused by N-terminal fusion of the reporter is unlikely, and (iii) *SPPiDDR1* was not expressed under basal (control) conditions, our data indicate that UV-C irradiation triggers both transcriptional activation and retention of SPPiDDR1 proteins at the periphery of peroxisomes (alternatively re-localization from the peroxisome lumen).

### Dicot genomes contain several SPPiDDR paralogs

The protein BLAST searches with ORF3 did not yield one, but several hits for each species, suggesting the presence of several *SPPiDDR* genes in plant genomes (**Fig. 5A**). In *Arabidopsis thaliana*, two additional hits (besides SPPiDDR1) were obtained: At4G12735 and At4G12731 (hereafter referred to as SPPiDDR2 and SPPiDDR3, respectively) (**Fig. 5A**). Note that the corresponding loci lie in close proximity to each other on chromosome 4 (only 642 bp separate the start codon of *At4G12731* from the stop codon of *At4G12735*) and are annotated as “hypothetical peptides” in Araport11. A SOG1-binding motif was found in the promoter of *SPPiDDR2,* like for *SPPiDDR1,* but not in the promoter of *SPPiDDR3*. On the amino acid level, all three SPPiDDRs are closely related: SPPiDDR1 shares 68% and 67% identity with SPPiDDR2 and SPPiDDR3, respectively; and the linked SPPiDDR2 and SPPiDDR3 sequences even share 75% identity. SPPiDDR2 and SPPiDDR3 are therefore the closest paralogs. In addition, SPPiDDR2 and SPPiDDR3 present all features previously described for SPPiDDR1: a conserved N-terminal end with the strictly conserved cysteine, a high content of serines, and a conserved C-terminal end with canonical PTS1 motif (**Fig. 5B**).

**Figure 5:**
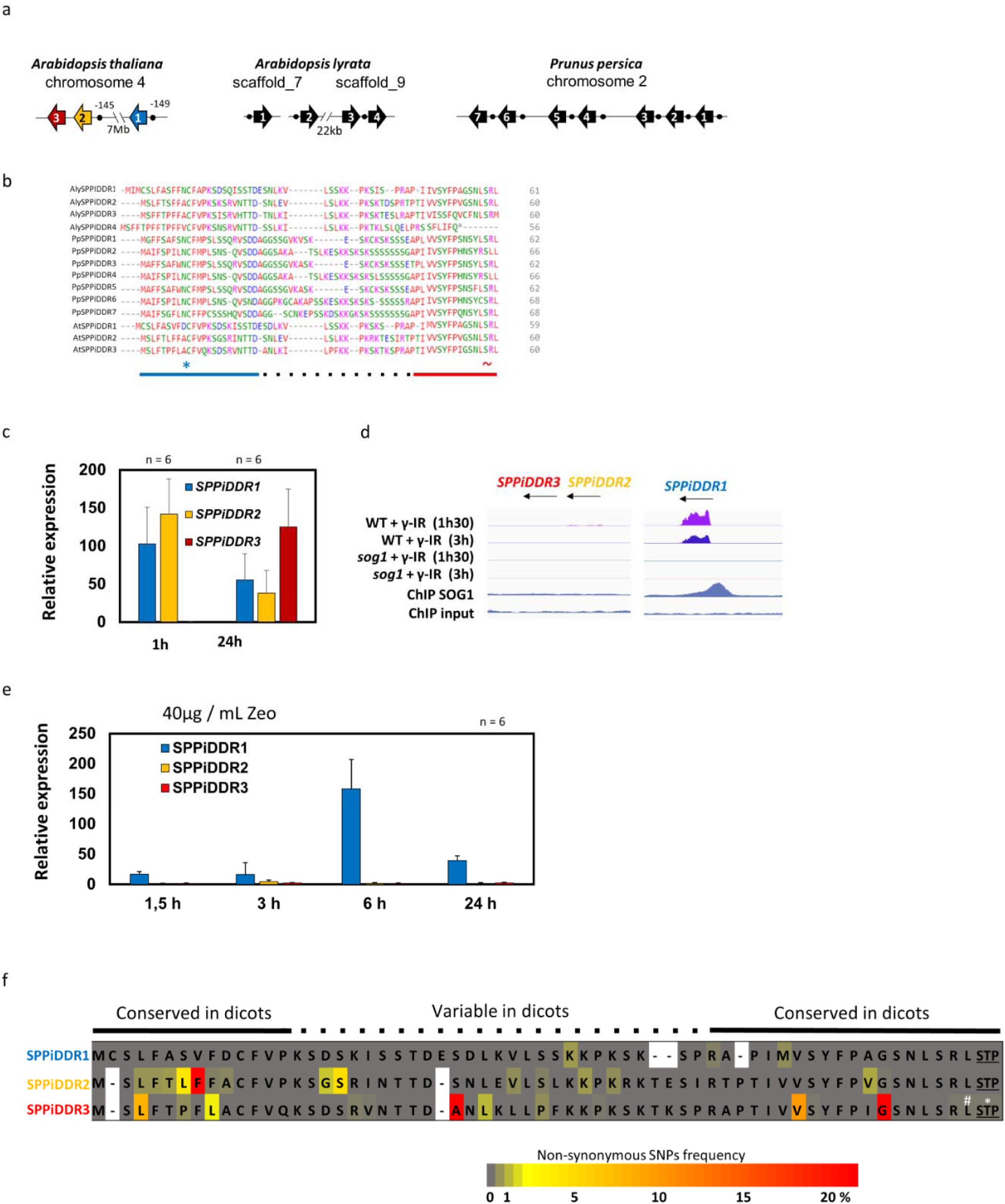
Multiple *SPPiDRR* copies coexist in dicots and have potentially diversified in *Arabidopsis thalinia*. a- Genomic representation of the 3 *SPPiDDRs* in *A. thaliana*, the 4 *SPPiDDRs* in *Arabidopsis lyrata*, and the 7 *SPPiDRRs* in *Prunus persic*a. Arrows indicate the orientation of the genes. Black dots are SOG1-binding motifs. b- Protein alignment of *SPPiDDRs* from panel (a). The blue line indicates the conserved N-terminal end with a strictly conserved cysteine (*), the red line indicates the conserved C-terminal end with the conserved SRL motif (∼), and the dotted line indicates the variable central area. c- Relative expression of *AtSPPiDDRs* 1 and 24 hours after UV-C irradiation (3 kJ/m²) compared to untreated plants. d- Examination with IVG software of *AtSPPiDDR* loci using data from Bourbousse et al., 2018. Reads from wild-type and *sog1* mutants 1 ½ hour or 3 hours after exposure or not (mock) to γ-IR (60 grays) were obtained with RNA-seq, reads of plants one 1 hour after exposure to γ-IR (60 grays) carrying a *pSOG1:SOG1-3xFLAG* construct as bait compared to their respective input controls were obtained from ChIP-Seq data. Scales are identical between RNA-seq reads and between ChIP-seq reads. Scales are identical between RNA-seq reads and between ChIP-seq. e- Transcription analysis over time by RT-qPCR of *SPPiDDR1*, *-2* and *-3* in response to zeocin (40 µg/mL). Data were normalized to the values in non-treated wild-type plants. f- Heat map diagram of a protein sequence alignment displaying the non-synonymous SNPs frequency (grey to yellow to red) in 1135 Arabidopsis accessions for the different Arabidopsis *SPPiDDR* paralogues. The white interruptions indicate gaps in the alignment. # indicates an earlier stop codon, and * indicates loss of the stop codon.

Even though information on these 2 new *Arabidopsis thaliana* loci in the scientific literature is scarce, data mining revealed that (i) *SPPiDDR2* is hardly expressed under basal conditions like *SPPiDDR1,* (ii) *SPPiDDR2* expression is strongly upregulated by 150 mM NaCl (1000 fold-change), submergence (220 fold-change), and to a lesser extent upon phosphate starvation or incubation with 0.1 µM cytokinin (4 and 5 fold-change, respectively); (iii) *SPPiDDR2* seems to be expressed predominantly in roots, leaves and in cell suspension, (iv) reporter-SPPiDDR2 fusions targeted peroxisomes when overexpressed in protoplasts, and (v) *SPPiDDR2* used to be annotated as non-coding RNA (see **Suppl. Fig. S3** and references^36–38^). In addition, by screening proteomic datasets of the scientific literature, we found peptides specific to SPPiDDR2 that were detected via mass spectrometry (MS) in Arabidopsis, confirming the existence of SPPiDDR proteins *in vivo*^39,40^. On the other hand, we found no study reporting on *SPPiDDR3*, which seems to be only weakly expressed (**Suppl. Fig. S3)**.

To assess whether the two additional *SPPiDDR* paralogs in *A. thaliana* were transcriptionally active, their expression was measured after UV-C irradiation (**Fig. 5C**). The *SPPiDDR2* expression kinetic was similar to *SPPiDDR1* with a strong induction after 1 hour and a decline after 24 hours. In contrast, *SPPiDRR3* was not induced after 1 hour but showed a strong induction after 24 hours, demonstrating that this locus can be activated too. The early UV-C induction seen with *SPPiDDR1* and *-2* correlates with the presence of a SOG1-binding motif in the promoter, while this motif is lacking in the *SPPiDDR3* promoter. By reanalyzing transcriptomic datasets of plants exposed to γ-IR^17^, we found that *SPPiDDR2* and *-3* were not induced at all, in contrast to *SPPiDDR1* (**Fig. 5D**). In line with this, no binding of SOG1 after γ-IR was found by ChIP-seq, neither to the *SPPiDDR2* nor the *SPPiDDR3* promoters. When we similarly monitored induction of *SPPiDDRs* upon zeocin treatment, only *SPPiDDR1* was induced, whereas *SPPiDDR2* and -*3* remained at basal expression levels (**Fig. 5E**). The same transcriptional pattern was found in response to X-rays when re-examining the transcriptomics data from Wang et al. (2016)^28^. Taken together, the data indicate that *SPPiDDRs* differ from each other transcriptionally, with *SPPiDDR1* being the most active paralog in response to known DNA damaging treatments.

We also examined in detail the genomes of two other species: *Arabidopsis lyrata* of the order of Brassicales and a close relative of *Arabidopsis thaliana*, as well as *Prunus persica*, a taxonomically more distant species that belongs to the Rosales order. Using nucleotide BLAST analyses, we found that the *Arabidopsis lyrata* genome contains 2 *SPPiDDR* genes (*AL7G15570* and *AL9U12080,* here renamed *AlySPPiDDR1* and *AlySPPiDDR2*) that were previously annotated in *Arabidopsis lyrata* v2.1. Moreover, we identified 2 non-annotated additional *SPPiDDR* genes forming a tandem on scaffold 9 (*AlySPPiDDR3* and *AlySPPiDDR4*) (**Fig. 5A**). A SOG1-binding motif was found in the promoter of *AlySPPiDDR1, -2,* and *-4*, but not *AlySPPiDDR3.* While AlySPPiDDR1, -2, and -3 proteins show high sequence identity to AtSPPiDDR1, AlySPPiDDR4 displays an early stop codon, leading to the loss of the last 8 amino acids, including the PTS1 motif (**Fig. 5B**). Moreover, in AlySPPiDDR3 the PTS1 has turned into a “SRM” motif, which likely still functions as a PTS1. In the *Prunus persica* genome, 7 *SPPiDDR* genes were found, all located side-by-side on the same locus and all containing ORFs with a high sequence identity to AtSPPIDDR1 (**Fig. 5A** and **5B**). A SOG1-binding motif was found in the promoter of each of the *PpSPPiDDR* genes, and all PpSPPiDDR proteins display a PTS1, although the canonical SRL was turned into SLL in two of them: PpSPPiDDR2 and -4. All in all, our data indicate that multiple *SPPiDDR* paralogs coexist within plant genomes. While they appear globally conserved, some *SPPiDDR* genes display nonetheless substantial differences.

### SPPiDDR orthologs are diverging while paralogs are co-evolving

To understand how the *SPPiDDR* paralogs and orthologs may have evolved, we created a protein identity tree with the 100 top sequences obtained by protein BLAST using the 15 C-terminal amino acids of AtSPPiDDR1 (**Suppl. Fig. S4**). AtSPPiDDR1, -2 and -3 grouped together with all other SPPiDDRs from the *Brassicaceae* species, whereas non-*Brassicaceae* dicot SPPiDDRs formed an independent clade, composed of smaller clusters with several SPPiDDRs from the same species, such as *Helianthus annuus*, whose 6 paralogs are very similar to each other. These data suggest that plant genomes tend to retain *SPPiDDRs* in multiple copies, with orthologs diverging from each other and paralogs evolving together. This last point suggests functional redundancy between co-evolving paralogs, rather than functional divergence and sub-functionalisation.

To comprehend the importance of *SPPiDDRs* in Arabidopsis, we analyzed the frequency of non-synonymous single nucleotide polymorphisms (SNPs) in the three paralogous *AtSPPiDDR* genes, considering >1000 Arabidopsis accessions aside of the reference accession Col-0. We found that *SPPiDDR1* was practically invariant with only 3 non-synonymous SNPs at very low frequency. The two other *SPPiDDR* paralogs were more versatile, with 15 non-synonymous SNPs each, including some whose frequency reached up to 20% (**Fig. 5F**). Furthermore, only *SPPiDDR3* displayed non-synonymous SNPs in the PTS1, an earlier stop codon or loss of the stop codon, albeit at extremely low frequency. Together with the fact that *SPPiDDR3* appears to be the least transcriptionally active of the Arabidopsis *SPPiDDRs* (**Fig. 5** and **Suppl. Fig. S4**), our data indicate that *SPPiDDR3* might be less important than *SPPiDDR1* and *SPPiDDR2*, if not already undergoing pseudogenization. *SPPiDDR2* seems less conserved than *SPPiDDR1* but is still transcriptionally active. It may therefore act redundantly with *SPPiDDR1*, or have sub-functionalized to some extent; alternatively, it might have started to get pseudogenized.

### SPPiDDR1 and -2 act redundantly, slowing down rosette growth under salt stress

It has been observed previously that the *SPPiDDR1* locus contributes to recovery following zeocin-induced DNA damage^30^. Yet, zeocin treatment only induced *SPPiDDR1*. In order to investigate simultaneously a potential redundancy between the *SPPiDDR*s and possible involvement in other stress responses, we tested a condition that induced two Arabidopsis paralogs. We found publicly available DNA insertion lines for *SPPiDDR1* (Gabi_276G08, referred as to *sppiddr1*) and *SPPiDDR2* (WiscDsLox461-464J23, referred as to *sppiddr2*), but none for *SPPiDDR3*. Then we searched for a condition that may induce both *SPPiDDR1* and *-2*. Using the *proSPPiDDR1::GFP-GUS* reporter line, we found that *SPPiDDR1* is induced by salt, like *SPPiDDR2* (**Suppl. Fig. S5)**^36^. Salt stress triggers a ROS burst that causes DNA oxidation, which in turn activates the SOG1 pathway, consequently linking salt stress and DDR^2^. As both *SPPiDDR1* and *-2* were induced by salt, we wondered whether these loci may be involved in salt-stress tolerance or sensitivity. We found that *SPPiDDR1* is expressed in all vegetative tissues, including roots (**Fig. 2B**), and also *SPPiDDR2* when considering independent transcriptomic datasets from other studies (**Suppl. Fig. S4**). Thus, NaCl (i) induces both *SPPiDDR1* and *-2*, (ii) indirectly causes DNA damage, and (iii) may easily be added as solution to water soil-grown plants, making salt treatment a convenient tool to search for *SPPiDDR*-related phenotypes.

We therefore scrutinized *sppiddr1* and *sppiddr2* single mutants upon watering with a 50 mM NaCl solution. Both *sog1* and *sos1* mutants were included as controls, because they are known to be hyper-tolerant and hyper-sensitive to salt stress, respectively^2,41^. To investigate a potential redundancy between the two paralogs, a *sppiddr1 sppiddr2* double mutant (hereafter referred as to *sppiddr1,2*) was generated by crossing. Impaired induction of the respective *SPPiDDRs* in *sppiddr1*; *sppiddr2*, and *sppiddr1,2* mutant plants was confirmed by qPCR (**Suppl. Fig. S6**). Interestingly, *SPPiDDR1* expression was reduced in the *sppiddr2* single mutant; and vice versa, *SPPiDDR2* expression was reduced in the *sppiddr1* single mutant. This finding might indicate that *SPPiDDR1* and *SPPiDDR2* promote each other’s transcriptional induction.

Under standard growth conditions in soil, both *sppiddr2* and the *sppiddr1,2* double mutant showed higher rosette weights than wild type and *sppiddr1* (**Fig. 6A** and **6B**). This suggests that *SPPiDDR2* attenuates rosette growth under normal conditions, which also indicates that in certain cells and/or developmental stages, *SPPiDDR2* is expressed despite the absence of stress. When grown under salt stress, *sppiddr1* and *sppiddr2* single mutants did not display any obvious phenotype, whereas the *sppiddr1,2* double mutant produced slightly bigger rosettes (**Fig. 6A**). This phenotype of *sppiddr1,2* became more obvious when considering the relative rosette weight, *i.e.* the weight of rosettes from plants watered with salt solution divided by that of plants watered normally (**Fig. 6C**). Apparently, the *sppiddr1,2* double mutant is less sensitive to salt stress than the wild type. In fact, this phenotype is reminiscent of *sog1* mutants, albeit not as pronounced. Together, our data indicate that *SPPiDDR1* and *2* confer salt-sensitivity by inhibiting plant growth in a redundant manner in the presence of salt. Furthermore, as the *sppiddr1,2* double mutant resembles the *sog1* mutant, *SPPiDDR1* and *-2* seem to play an important role in SOG1-dependent stress responses, at least at the growth level.

**Figure 6:**
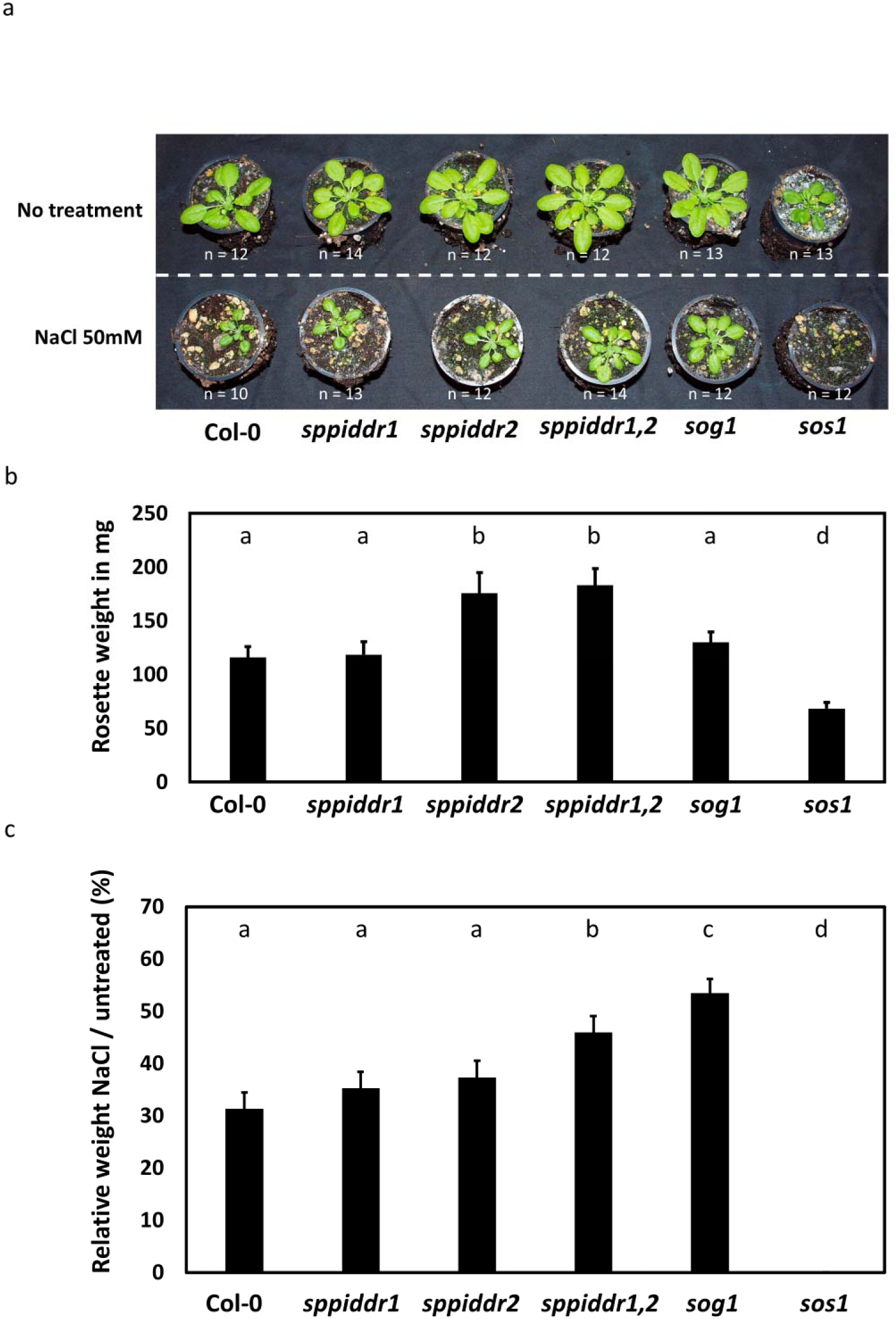
*SPPiDDR1* and *-2* act redundantly to slow down rosette growth under salt stress in Arabidopsis. a and b- Seeds were germinated and grown on soil watered every week with or without 50 mM NaCl. *sog1* and *sos1* mutants were included as partially insensitive and hypersensitive to salt stress, respectively. For (a), the picture was taken 4 weeks after germination. Then rosettes were decapitated and weighted. The values of the control condition (*i.e.* water) are shown in (b). c- The relative weights (*i.e.* weights with NaCl / weights without NaCl) from (b) are shown. The statistical analysis was performed via an ANOVA test. The data shown in these panels are representative of 3 independent experiments.

## DISCUSSION

### SPPiDDRs stood in a genomic and proteomic blind spot

LncRNAs are defined as loci with transcripts longer than 200 bases that do not encode proteins^42^. By selecting transcriptionally active loci that lack ORFs, bioinformatics-based studies have uncovered many lncRNAs in genomes of mammals and plants^43,44^. Yet, this approach suffers from several shortcomings. Many lncRNA are expressed only under specific conditions^45^ and may consequently remain unnoticed. Moreover, ruling out short ORFs may result in the overlooking of genes that code for small proteins. For instance, the Plant Long noncoding RNA Database (PLncDB) used a threshold of ORFs <100 amino acids, while the OFR threshold of the updated version (PLncDB V2.0) is even higher (120 amino acids)^31,32,36^. This implies that any transcriptionally active gene coding for a protein shorter than 120 or 100 amino acids is considered as lncRNA, a mis-annotation issue recently addressed by others^46^.

In this study we showed that the *RLSF_026432* locus is not a lncRNA, but a gene encoding a short peroxisomal protein of 59 amino acids that we named SPPiDDR1. Recently, several other alleged lncRNAs have also been found to be wrongly annotated^47–49^. In line with this, *SPPiDDR2* was previously also annotated as a non-coding sequence, but recently reported to encode a peroxisomal peptide^50^. By data mining, we identified peptides corresponding to SPPiDDR2 in proteomics databases of independent studies^39,40^, which proves that *SPPiDDRs* are translated *in vivo*. In both studies, peptides matching to SPPiDDR2 were found in membrane fractions, corroborating our observation that SPPiDDR1 tends to localize at the periphery of peroxisomes, at least when its expression was triggered by UV-C. Our data also indicate that *SPPiDDR1* is the most active paralog in Arabidopsis. Yet, no peptide specific to SPPiDDR1 was found in proteomic studies, which might be a result of the stress-specific expression and association with peroxisomal membranes, since most proteomic studies focused on cytosolic fractions. However, SPPiDDRs are likely soluble proteins and therefore rather membrane-associated, as overexpression of *SPPiDDR1* (in this study) and of *SPPiDDR2* in protoplasts^50^ in the absence of stress led to accumulation in the peroxisome matrix. Localization at the periphery therefore likely indicates interaction with peroxisomal membrane proteins, or co-expressed factors that attach to peroxisomes upon DDR. All in all, the mis-annotation of *SPPiDDR1*, the specific conditions of expression, plus the short length and peculiar subcellular localization of encoded proteins, placed the *SPPiDDRs* in a genomic and proteomic blind spot, which explains why this gene family has gone unnoticed up to now.

### The SOG1 pathway, and beyond?

Only two other studies have previously investigated the *SPPiDDR1* locus, and although the authors considered it as a lncRNA, some of our conclusions are concomitant. We previously described that *SPPiDDR1* expression is partially ATM-dependent in response to X-rays^28^, while Durut et al. (2023)^30^ showed that *SPPiDDR1* contributes to DDR. We also found that *SPPiDDR1* is induced and involved in DDR, and now extend these findings by uncovering a direct regulation of *SPPiDDR1* by SOG1, the master regulator of DDR in plants. This is in line with the partial ATM-dependent expression of *SPPiDDR1* that we previously found^28^, because SOG1 activation is controlled by ATM^10^. To examine the importance of *SPPiDDR1*, both Durut et al. (2023) and ourselves^30^ searched for conditions that may reveal a potential phenotype of the *sppiddr1* mutants. In our study, we used a T-DNA line with insertion at the 85^th^ base after the start codon of *SPPiDDR1* (line GABI_276G08), whereas Durut et al. (2023)^30^ created a CRISPR-Cas9 deletion mutant.

We now show that *SPPiDDR1* and *SPPiDDR2* act redundantly in response to salt stress and inhibit rosette growth. This redundancy is in line with the fact that both *SPPiDDR1* and *SPPiDDR2* are induced by salt stress. On the other hand, Durut et al. (2023)^30^ observed that seedlings of their deletion mutant were more sensitive to zeocin than wild-type plants. This is also consistent with our findings, as we observed that *SPPiDDR1* is the only paralog in Arabidopsis that is induced by zeocin, rolling out a potential redundancy between SPPiDDRs in response to this drug. However, it is intriguing that we observed a higher stress tolerance of the *sppiddr1,2* double mutant, while Durut et al. (2023)^30^ observed an oversensitivity of their *SPPiDDR1* CRISPR-Cas9 mutant. This is not necessarily a contradiction, as plants insensitive to DNA damage (such as *sog1* mutants) accumulate more DNA lesions than wild-type plants, but perform better in terms of growth^2,51^. In addition, Durut et al. (2023)^30^ and our study also converge regarding the conservation of *SPPiDDR1*. However, using the protein rather than the genomic sequence, we found conservation of this locus across all dicots, not only within the *Brassicaceae*. Together with the fact that plants lacking SPPiDDR1 accumulate more DNA-damage than wild-type plants^30^, our results indicate that SPPiDDR1 contributes to mediating the SOG1-orchestrated DDR. Nevertheless, we found that the truncated *SPPiDDR1* promoter that still contains the SOG1-binding site showed decreased activity, implying that elements other than the SOG1 binding motif probably contribute to the transcriptional activation. On the other hand, *SPPiDDR3* was found to be induced in response to UV-C despite the absence of a SOG1-binding motif, and it was reported recently that E2FB binds to the promoter of *SPPiDDR2*^52^. Thus clearly, *SPPiDDR* gene expression is not solely controlled by SOG1. Hence, investigating E2FB or other TFs for controlling *SPPiDDRs* will provide insight on whether these genes are only involved in DDR or beyond, such as oxidative stress responses.

### What are the cellular and biological functions of SPPiDDRs?

We found that SPPiDDR1 targets peroxisomes. Given the fact that the C-terminal PTS1 is conserved in SPPiDDR2 and SPPiDDR3, and that SPPiDDR2 labelled peroxisomes when overexpressed in protoplasts^50^, we hypothesize that all SPPiDDRs localize to these organelles. Not only do peroxisomes play a crucial role in ROS production^53^, but they also eliminate the majority of the hydrogen peroxide (H_2_O_2_) generated in response to *e.g.* salt stress by means of enzymes capable of detoxifying ROS, like catalases and peroxidases^54,55^. In fact, an excess of ROS or a deficient peroxisomal ROS-detoxifying system can result in damaged peroxisomes that are removed by autophagy to ensure normal operation of the cell, a process also called pexophagy^56,57^. Interestingly, while ATM is mostly known to act within the nucleus in response to DNA damage, it was reported to localize at peroxisomal membranes upon ROS accumulation to promote pexophagy in human cells, thus linking peroxisome homeostasis with the DDR^58,59^. By discovering a peroxisomal protein induced in response to DNA damage, we established for the first time a link between peroxisomes and the DDR in plant cells.

Proteins can be directed to peroxisomes via two pathways. Briefly, the PTS1 at the C-terminus of proteins is recognized in the cytosol and transported to peroxisomes by the shuttling receptor PEX5, which favors cargo with canonical PTS1 over others (weak PTS1)^59^. Proteins with N-terminal PTS2 are recognized by PEX7, which then interacts with PEX5 for joint transport to the peroxisome^60^. The PTS1 found in the AtSPPiDDRs is -SRL, one of the strongest PTS1 motifs. Whether SPPiDDRs prevent other proteins with weaker PTS1 from being delivered to peroxisomes, or they direct interacting proteins to peroxisomes has to be investigated by future studies. Alternatively, SPPiDDRs might also have a role inside peroxisomes and contribute to peroxisomal metabolism under stress.

### The AtSPPiDDR paralogs differ from each other

On the protein level, SPPiDDR1, -2 and -3 show high sequence identity with conserved features: a conserved cysteine in the N-terminal end, a high content in serine, and a C-terminal PTS1. However, they differ in their transcriptional profiles. This indicates that they may be controlled by different TFs. we found a SOG1-binding motif in the promoter of *SPPiDDR1* and *-2* but not *SPPiDDR3*. *SPPiDDR1* is induced by X-rays, UV-C, zeocin, γ-IR, bleomycin and salt, and we found that SOG1 controls its induction in response to UV-C^17,28,30^. However, *SPPiDDR1* induction was only partially impaired in *sog1* mutant plants, suggesting that (an)other TF(s) may also contribute to its transcriptional control. On the other hand, UV-C, salt, phosphate starvation, submergence, ozone and biotic stresses, like infection by Botrytis, induce *SPPiDDR2* transcription^36^ (**Suppl. Fig. S3**). Interestingly, we observed no transcriptional activation by zeocin, X-rays or γ-IR, all genotoxic stresses, despite presence of a SOG1-binding motif in its promoter. However, SOG1 ChIP-seq data were available only for γ-IR^17^, a treatment that does not induce *SPPiDDR2*. Hence, we cannot conclude on whether *SPPiDDR2* is also a direct target of SOG1, or not. Interestingly, the TF E2FB that governs a SOG1-independent pathway in DDR was found to bind to the *SPPiDDR2* promoter^52^. Whether *SPPiDDR1* is solely controlled by SOG1 and/or E2FB controls *SPPiDDR2* would help explain the transcriptional difference displayed by these two genes. Finally, *SPPiDDR3* was induced only by ozone and UV-C, albeit delayed compared to *SPPiDDR1* and *-2* (**Suppl. Fig. S3 and Fig. 5**). Clearly, the three *SPPiDDR* paralogs in Arabidopsis differ in their transcriptional regulation. Moreover, considering that SPPiDDR1 displays strong conservation within Arabidopsis accessions, in contrast to SPPiDDR2 and -3, it seems that SPPiDDR1 is the canonical paralog. As *SPPiDDR2* is active under normal conditions and acts redundantly with *SPPiDDR1* in response to salt stress, the gene is still functional and probably has either (partially) subfunctionalized or even neofunctionalized. On the other hand, the poor transcriptional activity and high frequency of non-synonymous SNPs hint at a pseudogenization of *SPPiDDR3*. In future experiments, double and triple *SPPiDDR* mutant combinations should be examined next to the single mutants to provide a better understanding of these questions. As *SPPiDDR2* and *SPPiDDR3* lie back-to-back on chromosome 4 and no mutant is publicly available for *SPPiDDR3*, this cannot be tackled by classical crossing and requires either deletion of the entire *SPPiDDR2::SPPiDDR3* genomic bloc using CRISPR-Cas technology, or a silencing approach.

Our data also show that the multiple *SPPiDDR* paralogs found in dicot species (*e.g. Prunus persica* or *Helianthus annuus*) are more similar to each other than to the *SPPiDDR* orthologs. Moreover, PpSPPiDDR2 and -4 carry a weaker PTS1 than SRL (SLL)^61^, potentially reflecting ongoing evolutionary divergence. In addition, an early stop codon in AlySPPiDDR4 suggests a weaker selection pressure than on the PTS1-containing Arabidopsis paralogs. This may indicate that the *SPPiDDR* loci constitute a hotspot for paralog evolution in dicots. Such a phenomenon has already been described for the *SUMO* gene family, and a similar approach based on phylogeny and co-linearity could help retrace the history and diversification of the *SPPiDDRs* in plant genomes^62^.

## MATERIALS AND METHODS

### Plant material and growth condition

The *Arabidopsis thaliana* ecotype Columbia-0 (Col-0) was used as wild type for all analyses. Seeds were vernalized for 2 days at 4°C and then grown on soil under long-day conditions (16 hours light / 8 hours darkness, 100 μE, 20°C, 60% humidity). Alternatively, *A. thaliana* seeds were surface-sterilized with 70% ethanol for 5 min, washed with autoclaved water twice, and sown on half-strength Murashige & Skoog (MS) medium (1⁄2 MS, 100 mM MES, pH 6.5)^63^ containing 0.8% (w/v) agar. After vernalization (2 days at 4°C), seeds were placed into short-day conditions (8 hours light / 16 hours darkness). The used *sog1-1* and *sos1* mutants were described before^9,41,64^; *sppiddr1* seeds were obtained from GABI-Kat (GABI_276G08) and *sppiddr2* seeds from NASC (SALK_032841).

### Cloning

*SPPiDDR1* flanking sequences were amplified by PCR with a high-fidelity Q5 polymerase (NEB) and primers combining recombination attB sites and the extremity of sequence. GGGGACCACTTTGTACAAGAAAGCTGGGTAGACCATTTAGATTGATAGCA and GGGGACAAGTTTGTACAAAAAAGCAGGCTCAAATCCTCTTCTTTAGGGTTT were used as primers to amplify and clone the intergenic sequence between *SPPiDDR1* and *AT4G35580*; and GGGGACCACTTTGTACAAGAAAGCTGGGTAATAGGAGTTTGAAAAACGTGT and GGGGACAAGTTTGTACAAAAAAGCAGGCTCAGTCTGAAAAGTGGTAATTGAATT TTG were used for intergenic sequence between *SPPiDDR1* and *AT4G35589*. Fragments with apparent length matching the expected ones (728 bp and 975 bp, respectively), were then extracted from the agarose gel and the DNA purified following the manufacturer instruction (NucleoSpin® Gel and PCR Clean-up kit (MACHEREY-NAGEL)). Each purified DNA fragment was inserted into *pDONR207* via gateway™ BP reactions (ThermoFisher), and then into *pKGWFS7* via gateway™ LR reactions (ThermoFisher), using *E.coli* DH10B cells and appropriate antibiotics (20 µg/mL gentamicin for vectors with *pDONR207* backbones, and 50 µg/mL spectinomycin for vectors with *pKGWFS7* backbones). For both steps, colony PCRs were first carried out with primers overlapping the DNA fragment and the backbone of the plasmid to ensure that the insert was correct. Then, the vectors were sequenced to confirm the cloned *proSPPiDDR1*. Each final vector (*i.e. SPPiDDR1* flanking sequences into *pKGWFS7*) was then introduced into *Agrobacterium tumefaciens* (strain GV3101) via electroporation. Agrobacteria with *pKGWFS7* backbone vectors were grown on plates and liquid cultures using 50 µg/mL spectinomycin and 15 µg/mL rifampicin.

The same procedure was applied for the truncated *SPPiDDR1* promoter, using the primers GGGGACAAGTTTGTACAAAAAAGCAGGCTCAGTCTGAAAAGTGGTAATTGAATT TTG and GGGGACAAGTTTGTACAAAAAAGCAGGCTCAGACCCTAATTTTAATCATA.

The vector *pKGWFS7* containing *proSPPiDDR1* was digested by HindIII and MfeI restriction enzymes (NEB) and then loaded on an agarose gel. The bigger fragment (10968 bp) was cut out and purified. A *proSPPiDDR1::ORF3-GFP-GUS* fragment (1314 bp) with ORF3 stop codon and GFP start codon deleted was synthesized (Invitrogen GeneArt Strings DNA Fragments - ThermoFischer), digested with HindIII and MfeI restriction enzymes and purified. The digested DNA fragment was then introduced into the digested and purified *pKGWFS7* backbone in order to obtain a *proSPPiDDR1::ORF3-GFP-GUS* cassette within *pKGWFS7*. This binary plasmid was then introduced into *Agrobacterium tumefaciens* (strain GV3101 carrying pMP90 binary vectors) via electroporation.

For expression in protoplasts, *ORF3* was PCR amplified from genomic DNA of Arabidopsis wild-type leaves using Sense Primer NNNACTAGTATGTGTTCTCTCTTTGCCTCC with Antisense Primer NNNGGATCCTTATAGACGCGAAAGG and cloned in frame behind the reporter gene via compatible restriction sites into p*m*GFP-NXn and pOFP-NXn vectors^65^.

### Plant or leaf transformations and microscopic analyses

Stable transformations of Arabidopsis were carried out by the floral dip method^66^. Positive transformants (T1) were selected on MS plates containing the selection marker kanamycin (50 μg/ml). Presence of the T-DNA was confirmed by PCR using a forward primer specific to *proSPPiDDR1* and a reverse primer specific for *GFP*. Confirmed transformants were transferred to soil for seed collection. The seeds of each transformed line (T2) were then sown on MS plates containing kanamycin (50 μg/ml). 10 seedlings from each line with a single insertion (survival rate of 75%) were transferred to soil for seed collection. 50 seeds (T3) of each T2 line were then sown on plates containing kanamycin, and T2 lines with a homozygous insertion of the expression cassette (*i.e.* 100% survival rate) were selected.

Transient expression in leaves of 4-week-old *Nicotiana benthamiana* plants was performed by Agro-infiltration according to methods described before^67^. Infiltrated plants were left for two days in the growth chamber, before being exposed to UV-C (3 kJ/m²) in a dark room. Control plants were placed into a darkroom without UV-C light for the same time.

Microscopic analyses were carried out on leaf discs collected 20 to 24 hours after UV-C exposure. Epifluorescence was analyzed with an Axioskop 2 plus microscope (Zeiss), and confocal microscopy with a Zeiss LSM equipped, both equipped with a GFP filter set (excitation: 488-490 nm; detection: 520 nm).

### Plant treatments

For chemical treatment, 21-day-old seedlings were transferred to liquid ½ MS medium with or without 1 μg/ml bleomycin or 40 μg/ml zeocin. For heat and cold stress, MS solid plates containing 21-day-old seedlings were placed in incubators at 35°C or 4°C for 3 hours, respectively, while keeping comparable illumination. For UV irradiation, the seedlings grown on ½ MS plates were placed in a darkroom and exposed to 3 kJ/m² (unless specified differently) dose of UV-C (G30T8 30W Germicidal Lamp with Emission at 255 nm), or a 2 kJ/m² of UV-A or UV-B. The dose was measured with UVTOUCH (sglux, DE). Controls were placed in a darkroom with no UV light. The plates with seedlings were then placed back in the growth chamber under short-day conditions and sampled for analysis after appropriate time.

### GUS staining

Histochemical GUS assays were performed according to previously published protocols^68^. Briefly, Arabidopsis seedlings were grown on soil. After 10 days, they were extracted from the soil without damaging the roots and transferred into water in 24-well plates. After 1 day of acclimation, they were exposed to UV-C light (3 kJ/m²) and returned to the growth chamber. 24 hours later, the seedlings were placed in the reaction buffer, vacuum infiltrated, and incubated at 37°C overnight. Subsequently, the samples were bleached with 70% ethanol for at least 4 hours at 37°C. Afterwards, they were placed in 50% glycerol for 1 hour each and then mounted on glass slides for microscopy.

### RNA isolation, cDNA synthesis and qPCR

The RNA isolation protocol used was performed as described previously^69^. qPCR reaction mixtures were set up according to the manufacturer’s instructions (Blue S’Green, Biozym, DE) and reactions were carried out on a CFX96™ Touch thermocycler (Bio-Rad, CA, USA). As a control for contamination with genomic DNA, two primers specific for the AGAMOUS-LIKE 68 intron (AGL68) were used. Only samples with a cycle threshold (Ct) value for AGL68L>L30 were used for further analysis. Moreover, by using two primer pairs specific for either the 5′ or the 3′ end of glyceraldehyde-3-phosphate-dehydrogenase (GAPDH), the integrity of the cDNA was determined. If the difference in Ct value of GAPDH5′ and GAPDH3′ was above 1.5, samples were excluded from further analysis. UBIQUITIN 10 (UBI) and ACTIN 2 (ACT2) were used as reference genes for normalization.

### Protoplast transfection and microscopic analyses

Isolation and transfection of Arabidopsis leaf mesophyll protoplasts was performed via a PEG based method, as described previously^65^. For co-expression of GFP-ORF3 or OFP-ORF3 with peroxisomal marker constructs, 5 µg of plasmid DNA were mixed prior to transfection - either with 5 µg of a luminal peroxisome marker: the last 50 amino acids of PGL3 (At5g24400) with PTS1 motif -SRL fused N-terminally to GFP, or 7 µg of an ER/peroxisomal membrane marker: PEX16 (At2g45690) fused C-terminally to OFP^35^. All constructs were driven by the CaMV-35S promotor (*pro35S*). The transfected protoplasts were kept at room temperature in the dark and analyzed after approximately 24-48 h using a confocal laser-scanning microscope (Leica TCS SP5). For GFP, the excitation wavelength was set to 488 nm and for OFP to 561 nm, with emissions detected in the range of 490-520 nm (GFP) or 590-620 nm (OFP).

### Bioinformatic analyses

ORF search was carried out with ORFfinder (www.ncbi.nlm.nih.gov/orffinder/). Nucleotide and protein BLASTs were carried out on https://blast.ncbi.nlm.nih.gov/Blast.cgi, restricting searches in green plants, in Brassicaceae, or in green plants excluding Brassicacaea using the “organisms” section. Parameters for nucleotide BLASTs: expected threshold = 0.5, word size = 28, match/mismatch score = 1/-2, gap cost = linear; for protein BLAST: expected threshold = 0.5, word size = 5, matrix = BLOSUM62, gap cost = existence 11 / extension 1. Queries were *AtSPPiDDR1* for nucleotide BLAST, and either AtSPPiDDR1 or the 15 C-terminal amino acids of AtSPPiDDR1 protein for protein BLASTs. Sequences were collected and used for multiple sequence alignments and protein-identity tree with Clustal Omega (https://www.ebi.ac.uk/Tools/msa/clustalo/).

Genomic analyses of *SPPiDDRs* in *Arabidopsis thaliana*, *Arabidopsis lyrata* and *Prunus persica* were carried out on Phytozome v13 (https://phytozome-next.jgi.doe.gov/). Homologous sequences to *AtSPPiDDR1* were identified using the nucleotide BLAST tool (BLASTN, expected threshold = -1, Matrix = BLOSUM62, word lenght = default), and localized using the JBrowse tool. SOG1-binding motif (CTT(N)_7_AAG) was searched by using SnapGene® software (from Dotmatics; available at snapgene.com).

For SNP analyses, sequence conservation among *AtSPPiDDR2* and *AtSPPiDDR3* genes was assessed using 1135 Arabidopsis accessions from POLYMORPH 1001 (https://tools.1001genomes.org/polymorph/index.html) and the sequence conservation of *AtSPPiDDR1* was assessed using the accession from the Salk *Arabidopsis thaliana* 1001 Genomes (http://signal.salk.edu/atg1001/3.0/gebrowser.php). The percentage of accessions that contained an amino acid other than the prevalent residue was determined for each position for the 3 Arabidopsis SPPiDDR paralogs. A heat map of the percentage of accessions containing a different residue at a particular position is depicted on the multiple sequence alignment of the AtSPPiDDRs.

### Plant crossing and phenotyping

Single *sppiddr1* and *sppiddr2* mutant plants were crossed to obtain *sppiddr1,2* double mutant lines. The segregating offspring obtained from F1 plants were genotyped by PCR using either two primers flanking the T-DNA insertion (for amplifying wild-type alleles): CATAAAATTAAAGCCTGCGCATA and TGGAAAACTTGATTGGGTGAG for *SPPiDDR1*, TCAATAGTATCAGCCTCATATATTTTCTTG and CAAGGATGTCTCTCTTCACGCTCT for *SPPiDDR2*, or one primer combined with a primer annealing to the T-DNA border (for amplifying mutant alleles): CATAAAATTAAAGCCTGCGCATA and ATAATAACGCTGCGGACATCTACATTTT for *sppiddr1*, TCAATAGTATCAGCCTCATATATTTTCTTG and AATAGCCTTTACTTGAGTTGGCGTAAAAG for *sppiddr2*. F2 plants homozygous for both *sppiddr1* and *sppiddr2* mutant alleles were selected for phenotyping.

For the phenotyping analysis, seeds were vernalized (2 days at 4°C), sown on tap water-soaked soil previously treated with the fungicide Previcur (1 ml/L) and grown in short-day conditions (8 hours light / 16 hours darkness, 20°C). Seedlings were watered every 5-7 days. For salt stress, tap water supplemented with 50 mM NaCl was used to soak the soil and to water the seedlings. After 4 weeks, plants were photographed and then decapitated. Rosettes were weighted to assess the fresh weight.

### Statistical analysis

Assumptions for normality and homogeneity of variance were tested by the Shapiro and Levene Test, respectively. Depending on the outcome of the assumption analysis, either an analysis of variance (ANOVA), Kruskal-Wallis, Wilcox Rank Sum or Student’s t-test was performed. Bonferroni or Sidak correction was used for P-value adjustment. The significance level was set to P < 0.05.

### Re-analysis of RNA-seq and ChIP seq data

RNA-seq and ChIP-seq data from plants treated with γ-IR were obtained from Bourbousse et al. (2018), accession no. GSE112773 in the Gene Expression Omnibus (GEO) database (https://www.ncbi.nlm.nih.gov/geo). Reads were depicted using the IVG software^70^. Scales of each track depicting either RNA-seq reads or ChIP-seq on the same figure are identical.

### Accession numbers

SPPiDDR1: At4G00215; SPPiDDR2: At4G12735; SPPiDDR3: At4G12731; SOG1: AT1G25580; PEX5: AT5G56290; PEX7: AT1G29260; ATM: AT3G48190; ATR: AT5G40820; LINDA: AT3G00800; GAPDH: AT3G26650; UBI10: AT4G05320; ACT2: AT3G18780; AGL68: AT5G65080; BRCA1: AT4G21070; CYCB1: AT4G37490; SMR7: AT3G27630; PLA2A: AT2G26560; E2FB: AT5G22220.

## AUTHOR CONTRIBUTIONS

VH and RK designed the research; EG, JH and VH performed transcriptional analyses; VH performed the examination of publicly available RNA-seq, ChIP-seq and ribosome profiling datasets on the *SPPiDDR* loci; VH performed cloning, transient and stable transformations in plants, microscopy on leaves, and GUS analysis on seedlings; AS and LL performed cloning, transformation of protoplasts and microscopy; VH performed BLAST analyses, sequence alignments and the protein identity tree, and variability in Arabidopsis accessions; VH, EG and JH performed phenotyping analyses; VH wrote the paper with inputs from all the authors.

## ACKNOWLEDGEMENTS

We thank Antonia Schmitt, Felicia Rauch, Melika Arast, Luise Pfannebecker, Carla Castro and Maria Singer for their practical assistance. We express our gratitude to all colleagues who supplied material and contributed to discussions during lab-meetings. This work was supported by a grant from the Deutsche Forschungsgemeinschaft (DFG KU715/12-1) to Reinhard Kunze.

## SUPPORTING INFORMATION

Supplemental Table S1. Protein and nucleotide BLAST analyses.

Supplemental Table S2. Primers used for qPCR analyses.

Supplemental Figure S1. Ribosomes bind DNA at the *RLFS_026432 ORF3* locus.

Supplemental Figure S2. *m*Cherry-GFP also localized at the periphery of peroxisomes.

Supplemental Figure S3. *SPPiDDR2* is induced by numerous abiotic stresses.

Supplemental Figure S4. Protein identity tree suggesting co-evolution of SPPiDDRs paralogs.

Supplemental Figure S5. Salt induces *SPPiDDR1* expression.

Supplemental Figure S6. *SPPiDDR1* and *SPPiDDR2* expression is impaired in the *sppiddr1,2* double mutant.

