## Supplemental Figures for "*SPPiDDRs*: a new gene family in dicot plants involved in DNA-Damage Response"

### Slide 1
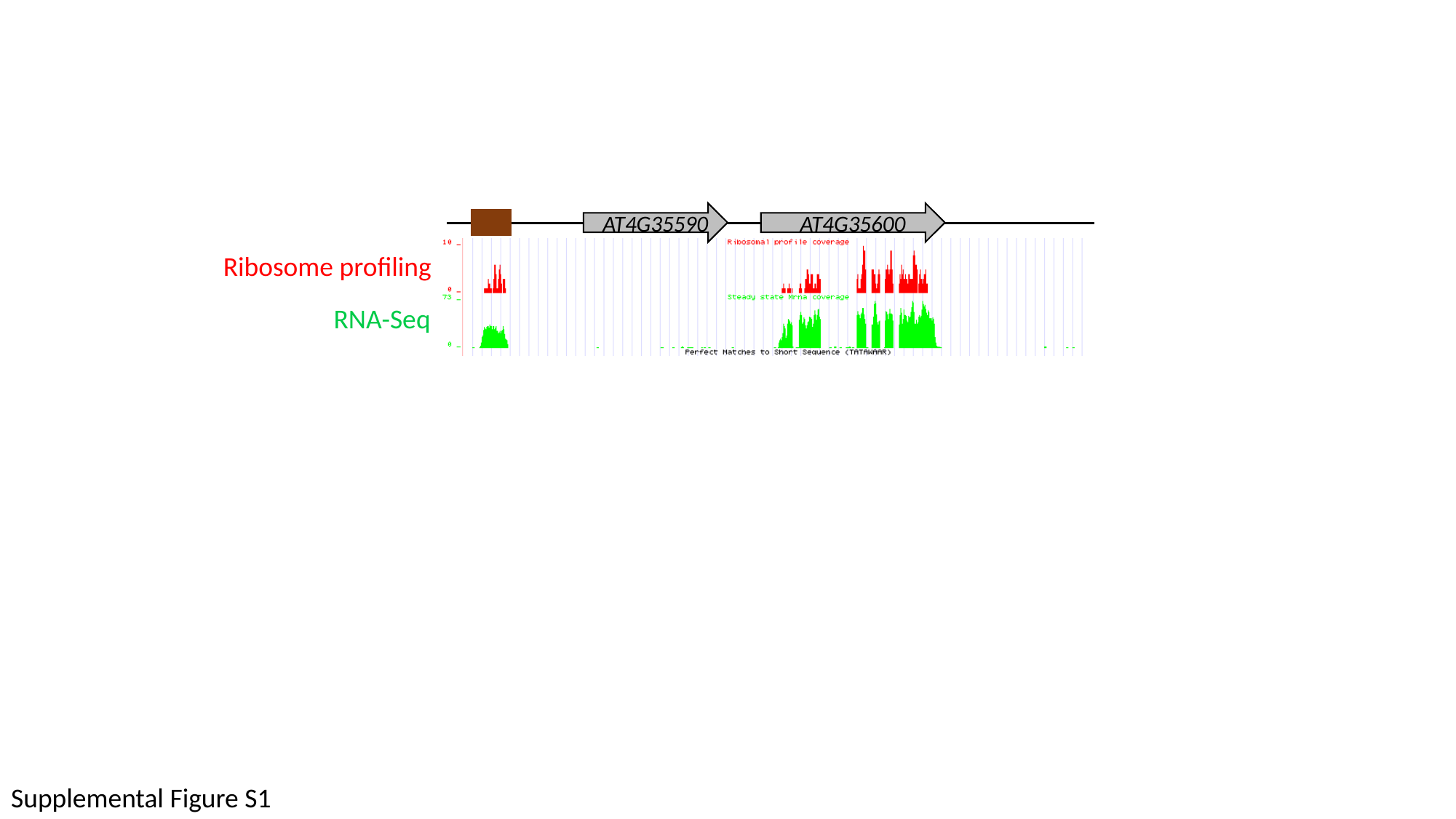

AT4G35590
AT4G35600
Ribosome profiling
RNA-Seq
Supplemental Figure S1

### Slide 2
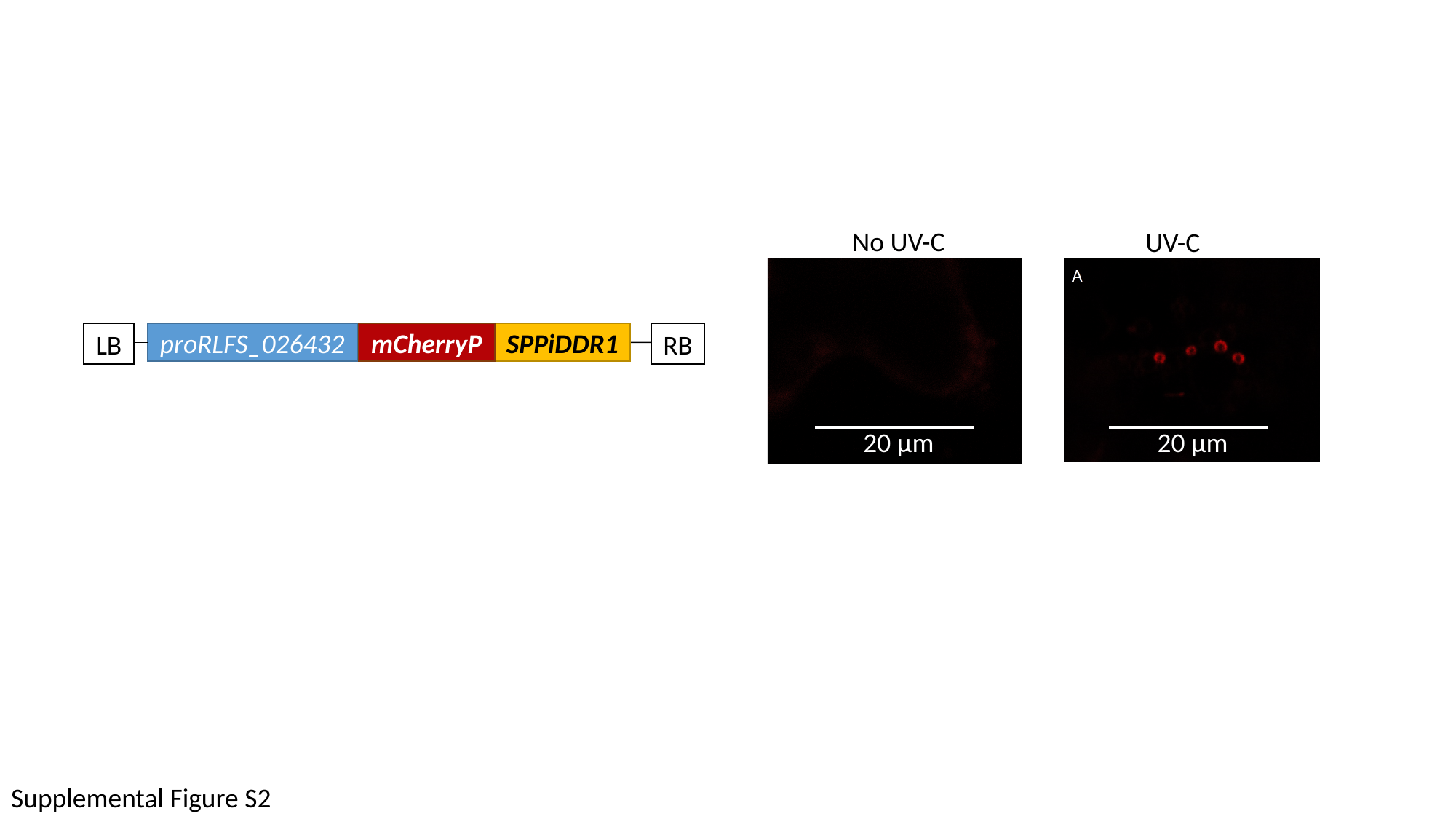

No UV-C
UV-C
SPPiDDR1
LB
proRLFS_026432
mCherryP
RB
20 μm
20 μm
Supplemental Figure S2

### Slide 3
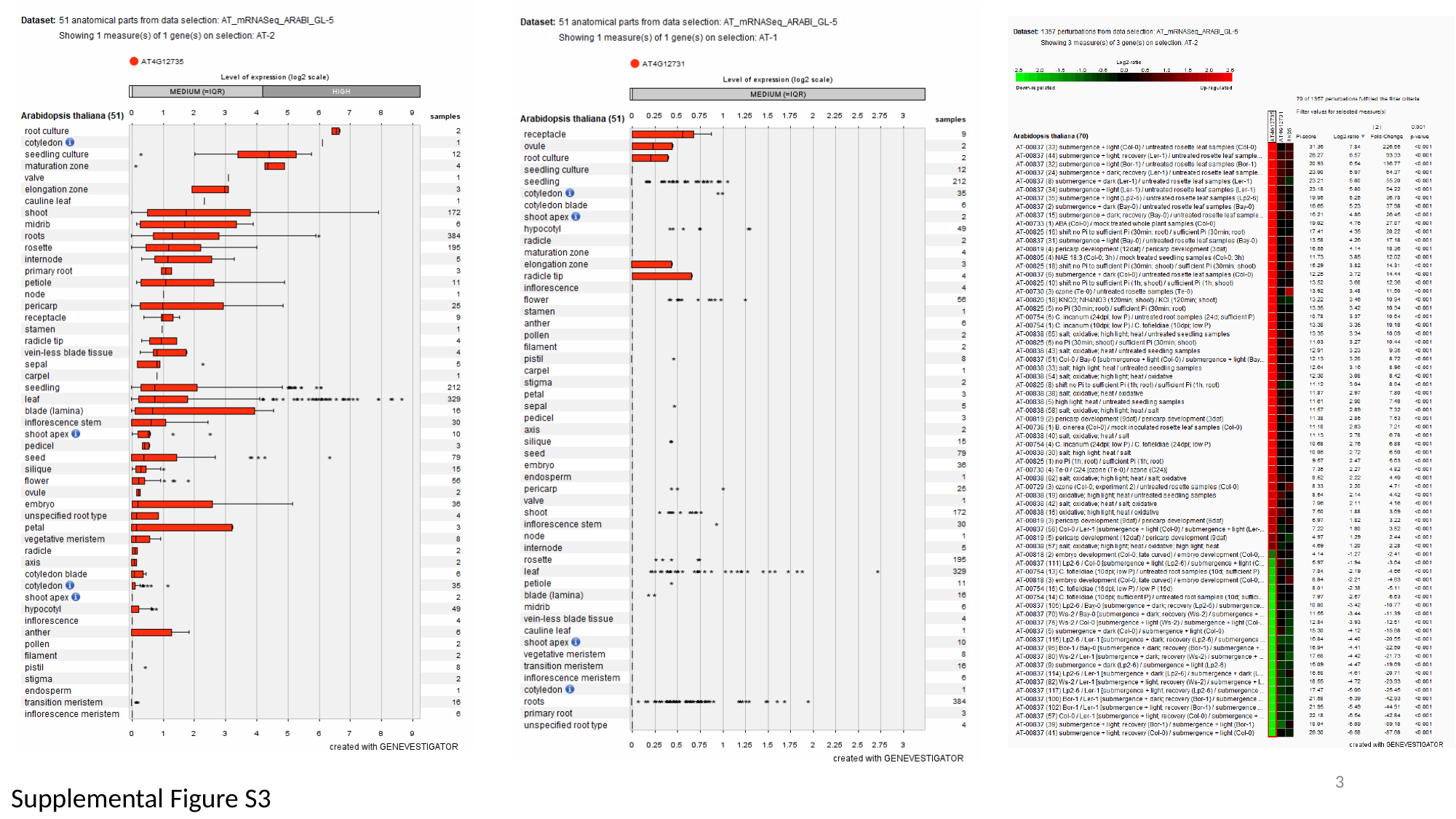

‹#›
Supplemental Figure S3

### Slide 4
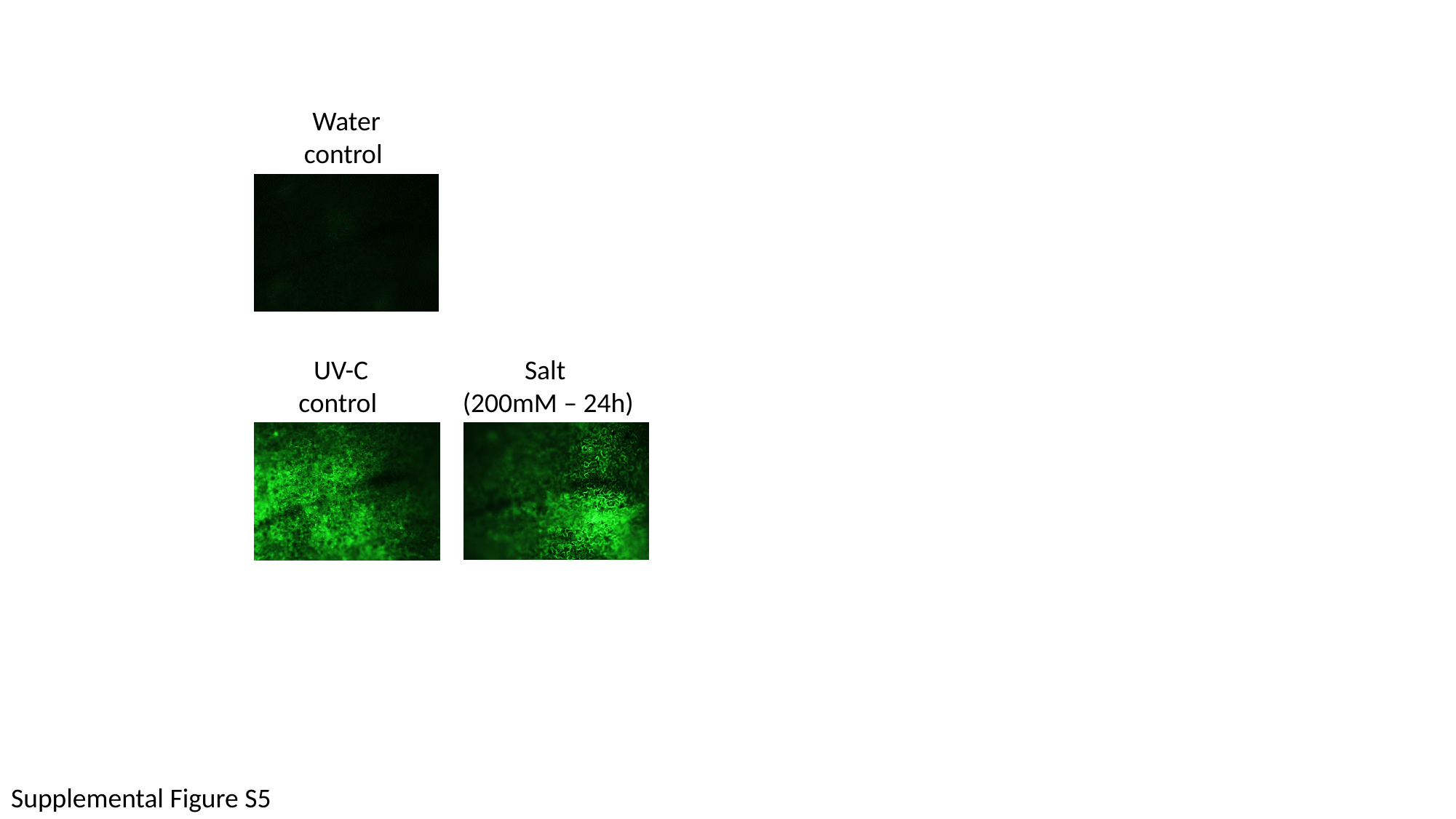

Water control
Salt
(200mM – 24h)
UV-C control
Supplemental Figure S5

### Slide 5
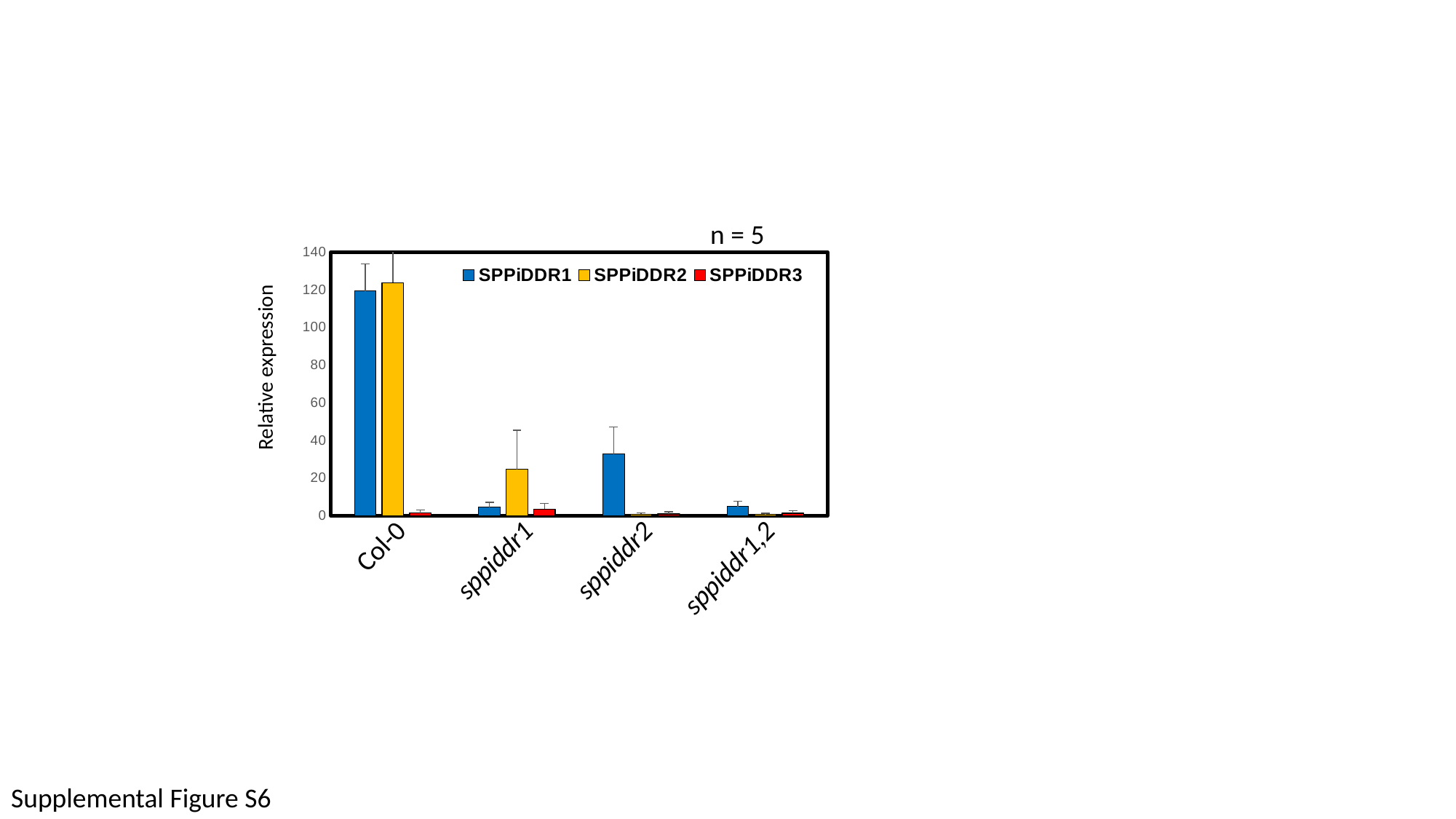

#### Chart
| Category | | | |
|---|---|---|---|n = 5
Relative expression
Col-0
sppiddr1
sppiddr2
sppiddr1,2
Supplemental Figure S6
