## Supplemental Information for "*SPPiDDRs*: a new gene family in dicot plants involved in DNA-Damage Response"

**Supplemental Materials & Methods**

*Ribosome-profiling data*

The transcriptomics and ribosome-profiling data are originally from Liu et al. (2013) and visualized here using the online server GWIPS-viz  (https://gwips.ucc.ie/).

Liu, M. J., Wu, S. H., Wu, J. F., Lin, W. D., Wu, Y. C., Tsai, T. Y., Tsai, H. L., & Wu, S. H.  Translational landscape of photomorphogenic Arabidopsis. *The Plant cell* **25(10)**, 3699–3710. doi.org/10.1105/tpc.113.114769 (2013)

*Cloning mcherry-SPP1DDR1 for confocal microscopy*

*pKGWFS7* containing the *proSPPiDDR1::SPPiDDR1-GFP-GUS* expression cassette was digested by HindIII and NcoI restriction enzymes and then loaded on an agarose gel. The bigger fragment (9389 bp) was cut out and purified. A *proSPPiDDR1::mcherry-SPPiDDR1* fragment flanked by HindIII and NcoI restriction site (1162 bp) was synthesized (Invitrogen GeneArt Strings DNA Fragments - ThermoFischer), digested with HindIII and NcoI  restriction enzymes and purified too. This digested DNA fragment was then introduced into the digested and purified pKGWFS7, in order to obtain a *proSPPiDDR1::mcherry-SPPiDDR1* cassette in pKGWFS7. This vector was then introduced into *Agrobacterium tumefaciens* (strain GV3101 carrying pMP90 binary vectors) via electroporation, which were then used for transient expression in tobacco leaves and subjected to microscopic analysis.

Microscopic analyses were carried out on leaf discs picked up 20 to 24 hours after UV-C exposure. Confocal microscopy was analyzed with a Zeiss LSM. The excitation wavelength was set to 587 nm and the emissions detected in the range of 595-625 nm

*Gene expression*

Tissue-specific expression patterns and condition-related expressions for *SPPiDDR2* and *SPPiDDR3* were obtained from Genevestigator ([www.genevestigator.com](http://www.genevestigator.com)). Data for *RWP-RK DOMAIN-CONTAINING 5* were added to the condition-related expressions for comparison.

*Protein-identity tree*

The protein sequences obtained from the BLAST analysis with the 15 C-terminal amino acids of AtSPPiDDR were used to build a protein-identity tree using Clustal Omega (<https://www.ebi.ac.uk/Tools/msa/clustalo/>) with settings left in the default mode. No distance corrections were applied on the obtained Neighbour-joining tree, and the branches are displayed with real lengths.

*Salt treatment on leaf-discs for SPPiDDR1 expression*

Plants were grown on soil for 4 weeks before excising leaf-discs, placed either in water (controls) or in water supplemented with NaCl (200 mM). As positive control for *SPPiDDR1* expression, some leaf-discs in water were exposed to UV-C (3 kJ/m²). The fluorescence was observed 24 hours later with an Axioskop 2 plus microscope (Zeiss) equipped with a GFP filter set (excitation : 488-490 nm; detection: 520 nm).
